# Rescaling protein-protein interactions improves Martini 3 for flexible proteins in solution

**DOI:** 10.1101/2023.05.29.542689

**Authors:** F. Emil Thomasen, Tórur Skaalum, Ashutosh Kumar, Sriraksha Srinivasan, Stefano Vanni, Kresten Lindorff-Larsen

## Abstract

Multidomain proteins with flexible linkers and disordered regions play important roles in many cellular processes, but characterizing their conformational ensembles is difficult. We have previously shown that the coarse-grained model, Martini 3, produces too compact ensembles in solution, that may in part be remedied by strengthening protein–water interactions. Here, we show that decreasing the strength of protein–protein interactions leads to improved agreement with experimental data on a wide set of systems. We show that the ‘symmetry’ between rescaling protein–water and protein–protein interactions breaks down when studying interactions with or within membranes; rescaling protein-protein interactions better preserves the binding specificity of proteins with lipid membranes, whereas rescaling protein-water interactions preserves oligomerization of transmembrane helices. We conclude that decreasing the strength of protein–protein interactions improves the accuracy of Martini 3 for IDPs and multidomain proteins, both in solution and in the presence of a lipid membrane.

## Introduction

Intrinsically disordered proteins (IDPs), folded proteins with long disordered tails, and multidomain proteins with folded domains connected by flexible linkers, are characterized by their high level of conformational dynamics. Molecular dynamics (MD) simulations provide a valuable tool for studying IDPs and multidomain proteins, as they can be used to determine full conformational ensembles at atomic resolution (***Thomasen and Lindorff-Larsen, 2022***). However, there are two central challenges that must be overcome for MD simulations to provide a useful description of such systems: the force field describing all the bonded and non-bonded interactions between atoms in the system must be sufficiently accurate *and* the conformational space of the protein must be sufficiently sampled (***Bottaro and Lindorff-Larsen, 2018***).

One way to address the challenge of sufficient sampling is to use coarse-grained (CG) MD simulations in which groups of atoms are represented as single beads (***Ingólfsson et al., 2014***). Martini is a widely used CG model in which 2–4 non-hydrogen atoms are represented by a single bead (***Marrink et al., 2007; Monticelli et al., 2008***). An attractive aspect of Martini is its modular structure and high degree of transferability, which allows the simulation of complex systems containing several different classes of biomolecules. The current version of Martini, Martini 3, shows improvements over previous versions in areas such as molecular packing, transmembrane helix interactions, protein aggregation, and DNA base pairing (***Souza et al., 2021***).

We have previously shown that Martini 3 simulations of IDPs produce overly compact conformational ensembles, resulting in poor agreement with small-angle X-ray scattering (SAXS) and paramagnetic relaxation enhancement (PRE) experiments (***Thomasen et al., 2022***). Using an approach inspired by previous work on assessing and rebalancing non-bonded interactions in Martini (***Stark et al., 2013; Javanainen et al., 2017; Berg et al., 2018; Berg and Peter, 2019; Alessandri et al., 2019; Larsen et al., 2020; Benayad et al., 2021; Majumder and Straub, 2021; Lamprakis et al., 2021; Martin et al., 2021***) and atomistic force fields (***Best et al., 2014***), we found that agreement with SAXS and PRE data could be significantly improved by uniformly increasing the strength of non-bonded Lennard-Jones interactions between protein and water beads by ~10% (***Thomasen et al., 2022***). This was also shown to be the case for three multidomain proteins, hnRNPA1, hisSUMO-hnRNPA1, and TIA1; however, due to the small sample size and the similarity between these three proteins, it remains an open question whether the approach generalizes to other multidomain proteins.

Our previous work was concerned with the properties of proteins in aqueous solution in the absence of other classes of biomolecules. Intuitively, increasing the strength of protein-water interactions should affect the affinity between proteins and other biomolecules. As a prototypical example, one would expect that increasing protein-water interactions would decrease the affinity of proteins for lipid membranes, since the interaction is tuned by the relative affinity of proteins for water vs. the membrane environment. The extent to which our previously described force field modification affects protein-membrane interactions, however, remains unclear. There is increasing evidence that IDPs and disordered regions play important physiological roles at lipid membranes (***Kjaergaard and Kragelund, 2017; Zeno et al., 2018; Das and Eliezer, 2019; Fakhree et al., 2019; Cornish et al., 2020***), and so it is important to understand better how force field changes that improve the description of disordered proteins in solution affect their interactions with membranes. In this context, it is important to note that unmodified Martini 3 has been quite successful at reproducing the specific membrane interactions for peripheral membrane proteins, as we previously showed (***Srinivasan et al., 2021***).

For previous versions of Martini, problems with overestimated protein-protein interactions have been corrected either by increasing the strength of protein-water interactions (***Berg et al., 2018; Berg and Peter, 2019; Larsen et al., 2020; Martin et al., 2021***) or by decreasing the strength of interactions between protein beads (***Stark et al., 2013; Javanainen et al., 2017; Benayad et al., 2021***). We hypothesize that for proteins in solution, the two force field corrections likely have similar effects, simply rebalancing the relative energies associated with hydration versus self-interaction. However, in the case of mixed systems, for example with proteins, water, and membranes, we might expect clearer differences between these approaches. For example, decreasing the strength of protein-protein interactions may better retain the affinity between proteins and other molecules as originally parameterized, while increased protein-water interactions may lower this affinity (Fig. 1). Thus, it remains an open question whether this specificity is retained when protein-water interactions are increased, and whether rescaling protein-protein interactions provides equivalent or improved agreement with experimental observations, both in comparison with unmodified Martini 3 and Martini 3 with rescaled protein-water interactions. We note that Martini 3 has already been shown to provide good agreement with free energies of dimerization for transmembrane proteins (***Souza et al., 2021***), so the major focus of this work is to rebalance the interactions of proteins in solution to improve the agreement with experiments.

**Figure 1.**
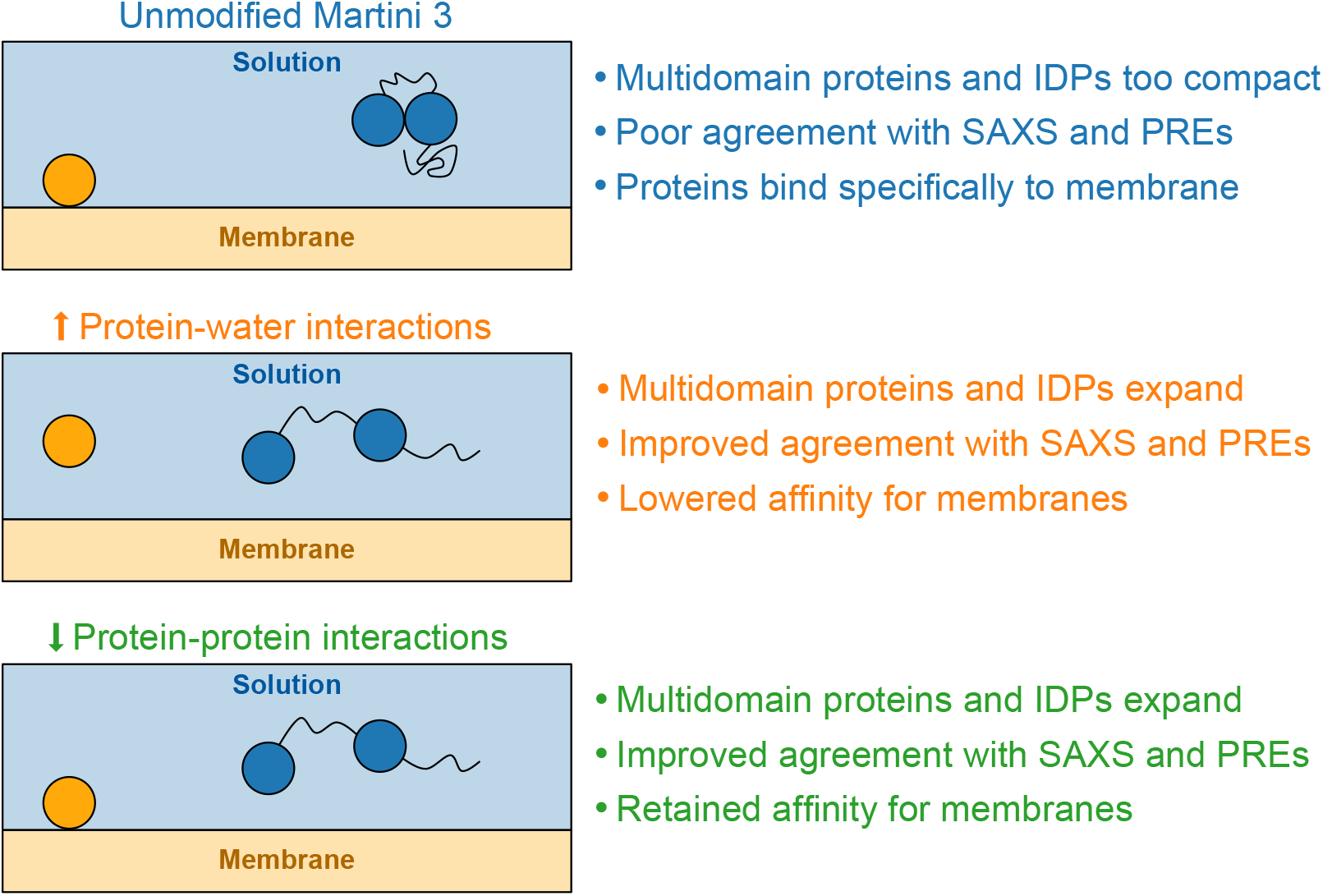
Expected effects of proposed force field modifications. Schematic overview showing the expected effects of rescaling protein-water and protein-protein interactions in Martini 3. Overestimated compactness of soluble IDPs and multidomain proteins and specific membrane interactions for peripheral membrane proteins have previously been reported (***Srinivasan et al., 2021; Thomasen et al., 2022***).

Here, we expand upon our previous work to address these questions. First, we have expanded the set of multidomain proteins to include 15 proteins for which SAXS data have previously been collected (Fig. 2). Using this five-times larger set of proteins, we show that, as was the case for IDPs, increasing the strength of protein-water interactions by 10% improves the agreement with SAXS data. We further show that decreasing the strength of non-bonded interactions between protein beads by 12% leads to a comparable improvement in agreement with SAXS and PRE data for IDPs and multidomain proteins in solution, but better preserves the specificity of protein-membrane interactions for peripheral membrane proteins. In contrast, we find that rescaling protein-protein interactions decreases the propensity of transmembrane helices to dimerize, whereas this propensity is mostly unchanged when rescaling protein-water interactions.

**Figure 2.**
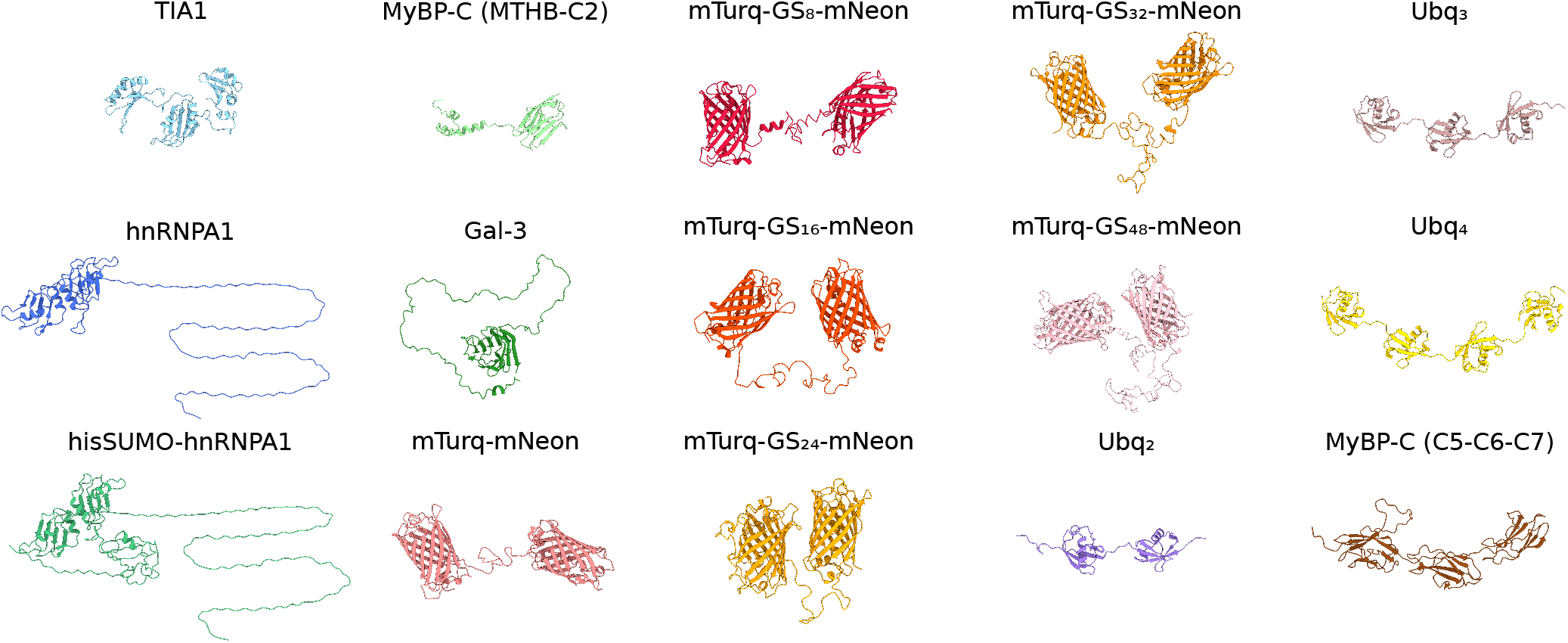
Starting structures for simulations of multidomain proteins. Starting structures of multidomain proteins used for Martini simulations. See the Methods section for a description of the source of the structures and how they were assembled.

## Results

### Analysis of an expanded set of multidomain proteins

Previously, we tested Martini 3 using a set of three multidomain proteins, TIA1, hnRNPA1, and hisSUMO-hnRNPA1, for which SAXS data have been measured (***Sonntag et al., 2017; Martin et al., 2021***). Given the similarity of the three proteins (all three are RNA-binding proteins, and the two latter differ only in the addition of a hisSUMO-tag), we wished to expand the set of proteins with mixed regions of order and disorder to include a wider range of sizes and domain architectures. We searched the literature for such proteins with reported SAXS data and identified 12 proteins that we added to our set (Fig. 2): the tri-helix bundle of the m-domain and the C2 domain of myosin-binding protein C (MyBP-C_MTHB-C2_) (***Michie et al., 2016***); the C5, C6, and C7 domains of myosin-binding protein C (MyBP-C_C5-C6-C7_) (***Nadvi et al., 2016***); linear ditri- and tetraubiquitin (Ubq_2_, Ubq_3_, Ubq_4_) (***Jussupow et al., 2020***); the two fluorescent proteins mTurquoise2 and mNeonGreen connected by a linker region with the insertion of 0, 8, 16, 24, 32, or 48 GS repeats (mTurq-GS_X_-mNeon) (***Moses et al., 2024***); and Galectin-3 (Gal-3) (***Lin et al., 2017***). Apart from Gal-3, these proteins all contain at least two distinct folded domains, connected by linkers of different lengths and composition; three proteins (Gal-3, hnRNPA1, and hisSUMO-hnRNPA1) also contain a long disordered region attached to a folded domain. Collectively, we will refer to this set as multidomain proteins, though we note that Gal-3 only contains a single folded domain.

We have previously shown that Martini 3 produces conformational ensembles that are more compact than found experimentally for a set of 12 IDPs and for the three mul-tidomain proteins TIA1, hnRNPA1, and hisSUMO-hnRNPA1, and that rescaling ε in the Lennard-Jones potential between all protein and water beads by a factor *λ*_PW_=1.10 resulted in more expanded ensembles that substantially improved the agreement with SAXS data (***Thomasen et al., 2022***). Using our much larger set of multidomain proteins, we examined whether Martini 3 generally produces too compact conformational ensembles of multidomain proteins, and whether our modified force field with rescaled protein-water interactions would generalize to the expanded set of proteins. We ran Martini 3 simulations of the 12 new multidomain proteins with unmodified Martini 3 and with *λ*_PW_=1.10 and calculated SAXS intensities from the simulations. We found that, on average across the 15 proteins, increasing the strength of protein-water interactions by *λ*_PW_=1.10 substantially improved the direct agreement with the experimental SAXS data, as quantified by the reduced *χ*^2^, 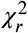 (Fig. 3). For only one of the 15 proteins, MyBP-C_MTHB-C2_, the modified force field gave rise to reduced agreement with the SAXS data. This result shows that our previously proposed modification of protein-water interactions in Martini 3, which was optimized to improve the global dimensions of IDPs, also provides a general improvement in the global dimensions of multidomain proteins.

**Figure 3.**
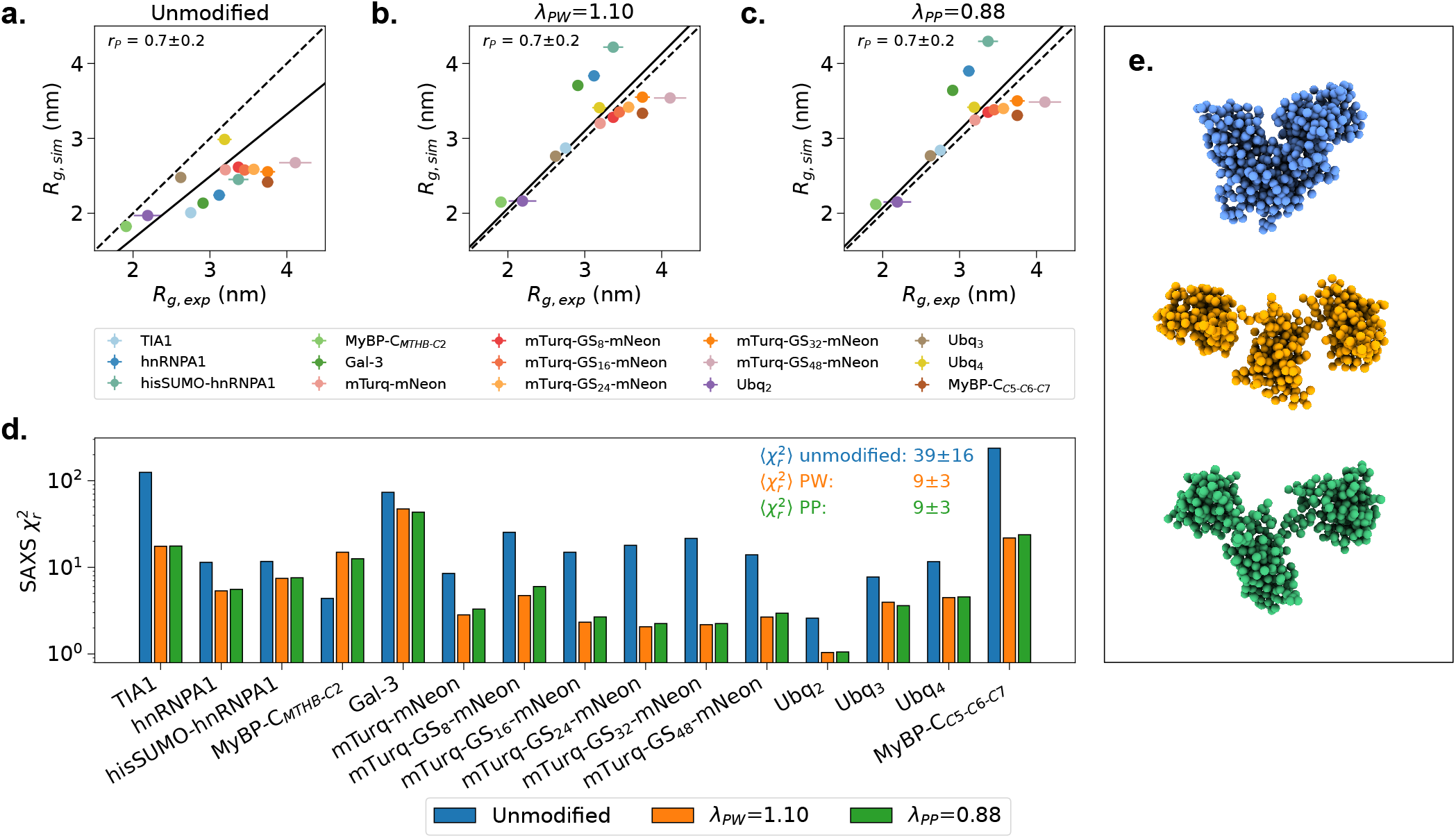
Agreement between simulations and SAXS data for multidomain proteins. *R*_*g*_ calculated from simulations plotted against *R*_*g*_ determined from Guinier fits to the SAXS data for **a** simulations with unmodified Martini 3, **b** simulations with protein-water interactions in Martini 3 rescaled by *λ*_PW_=1.10, and **c** simulations with protein-protein interactions in Martini 3 rescaled by *λ*_PP_=0.88. The diagonal is shown as a dashed line and a linear fit with intercept 0 weighted by experimental errors is shown as a solid line. Pearson correlation coefficients (*r*_*P*_) with standard errors from bootstrapping are shown on the plots. **d**. Reduced *χ* ^2^ to experimental SAXS intensities given by SAXS intensities calculated from unmodified Martini 3 simulations (blue) and Martini 3 simulations with protein-water interactions rescaled by *λ*_PW_=1.10 (orange) or protein-protein interactions rescaled by *λ*_PP_=0.88 (green). Mean and standard error of the mean over all proteins are shown on the plot. Note the logarithmic scale for 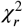. **e**. Representative conformation of TIA1 with an *R*_*g*_ corresponding to the average *R*_*g*_ in (blue) simulations with unmodified Martini 3, (orange) simulations with protein-water interactions in Martini 3 rescaled by *λ*_PW_=1.10, and (green) simulations with protein-protein interactions in Martini 3 rescaled by *λ*_PP_=0.88. Simulations of hnRNPA1, hisSUMO-hnRNPA1 and TIA1 with *J*_PW_=1.10 were taken from ***Thomasen et al. (2022***).

### Rescaling protein-protein interactions

Inspired by previous work on earlier versions of the Martini force field, (***Stark et al., 2013; Javanainen et al., 2017; Benayad et al., 2021***), we next examined whether rescaling protein-protein interactions instead of protein-water interactions would provide a similar or further improvement in the agreement with the experimental data. To do so, we ran Martini 3 simulations for the set of 12 IDPs with SAXS data available that we had studied previously (***Thomasen et al., 2022***) and the new set of 15 multidomain proteins. In these simulations, we rescaled ε in the Lennard-Jones potential between all protein beads by a factor *λ*_PP_. We scanned different values of this parameter, and found *λ*_PP_=0.88 to provide the best agreement with experiments (Fig. S1). We found that this level of rescaling protein-protein interactions (*λ*_PP_=0.88) provides a comparable improvement in the agreement with the experimental data as rescaling protein-water interactions by *λ*_PW_=1.10 for both multidomain proteins (Fig. 3) and IDPs (Fig. 4).

**Figure 4.**
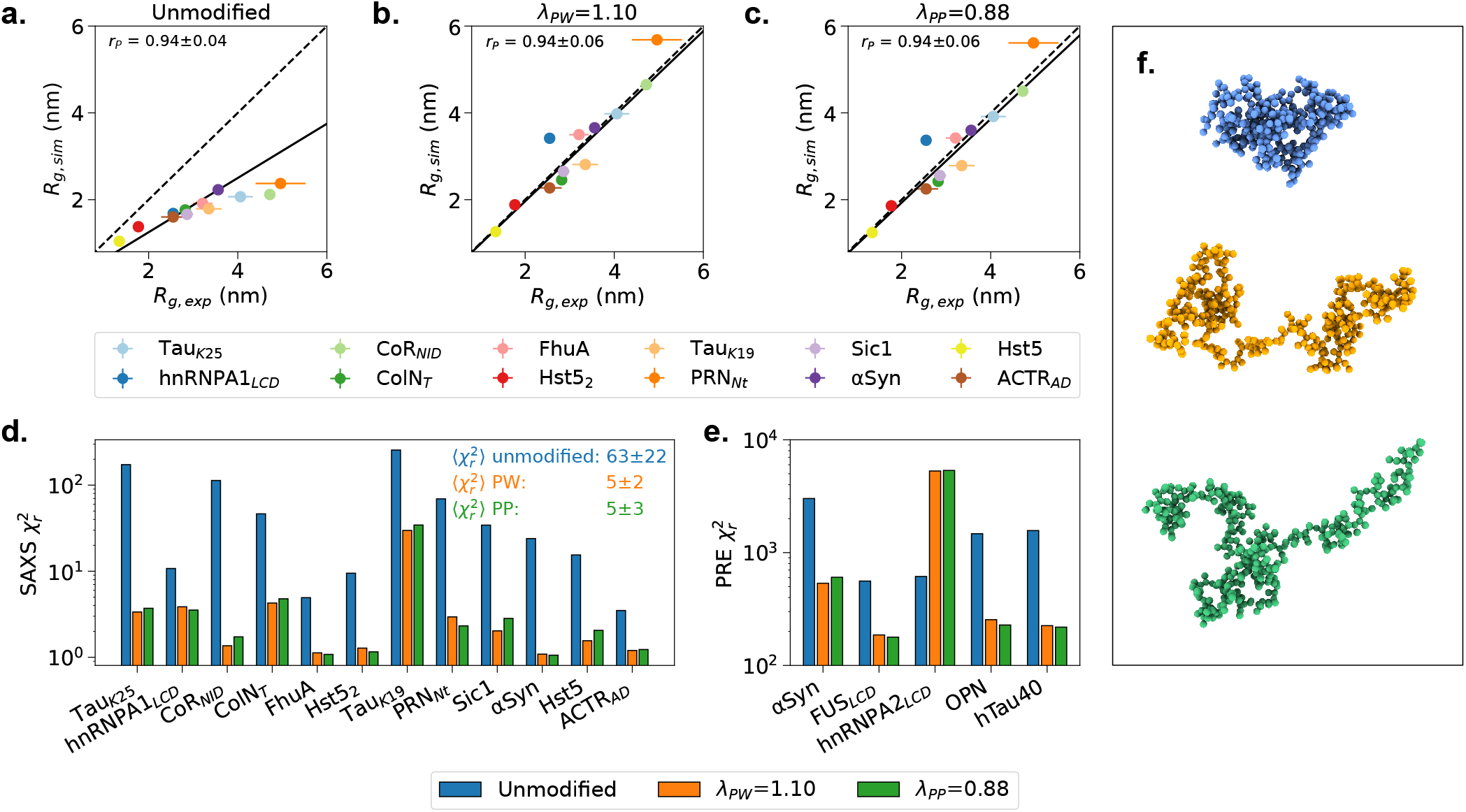
Agreement between simulations and SAXS or PRE data for IDPs. *R*_*g*_ calculated from simulations plotted against *R*_*g*_ determined from Guinier fits to the SAXS data for **a** simulations with unmodified Martini 3, **b** simulations with protein-water interactions in Martini 3 rescaled by *λ*_PW_=1.10, and **c** simulations with protein-protein interactions in Martini 3 rescaled by *λ*_PP_=0.88. The diagonal is shown as a dashed line and a linear fit with intercept 0 weighted by experimental errors is shown as a solid line. Pearson correlation coefficients (*r*_*P*_) with standard errors from bootstrapping are shown on the plots. **d**. Reduced *χ* ^2^ to experimental SAXS intensities given by SAXS intensities calculated from unmodified Martini 3 simulations (blue) and Martini 3 simulations with protein-water interactions rescaled by *λ*_PW_=1.10 (orange) or protein-protein interactions rescaled by *λ*_PP_=0.88 (green). Mean and standard error of the mean over all proteins are shown on the plot. Note the logarithmic scale for 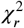. **e**. Reduced *χ* ^2^ to experimental PRE NMR data given by unmodified Martini 3 simulations (blue) and Martini 3 simulations with protein-water interactions rescaled by *λ*_PW_=1.10 (orange) or protein-protein interactions rescaled by *λ*_PP_=0.88 (green). Note the log-arithmic scale for 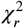. **f**. Representative conformation of Tau_K25_ with an *R*_*g*_ corresponding to the average *R*_*g*_ in (blue) simulations with unmodified Martini 3, (orange) simulations with protein-water interactions in Martini 3 rescaled by *λ*_PW_=1.10, and (green) simulations with protein-protein interactions in Martini 3 rescaled by *λ*_PP_=0.88. All simulations with *λv*_PW_=1.10 were taken from ***Thomasen et al. (2022***).

To further test the effect of rescaling protein-protein interactions by *λ*_PP_=0.88 and compare with the approach of rescaling protein-water interactions, we ran simulations of five IDPs with intramolecular PRE data available: the LCD of hnRNPA2 (***Ryan et al., 2018***), the LCD of FUS (***Monahan et al., 2017***), α-synuclein (***Dedmon et al., 2005***), full-length tau (hTau40) (***Mukrasch et al., 2009***), and osteopontin (OPN) (***Platzer et al., 2011***), and calculated PRE data from the simulations (Fig. 4e). Again, *λ*_PP_=0.88 provided a comparable level of agreement with the PRE data as we previously found using *λ*_PW_=1.10 (***Thomasen et al., 2022***). Specifically, the agreement with the PRE data improved for all proteins except the hnRNPA2 LCD (Fig. 4e).

To further characterize the symmetry between rescaling protein-water and protein-protein interactions, we compared the ensembles produced with the two rescaling approaches, and with unmodified Martini 3, using the distribution of *R*_*g*_ (Fig. S2-3) and a principal component analysis (PCA) based on the pairwise distances between backbone beads (Fig. S4-5). This analysis confirmed that the two rescaling approaches produce ensembles which are highly similar when compared with unmodified Martini 3. We conclude that decreasing the strength of protein-protein interactions by *λ*_PP_=0.88 provides an equally good alternative to rescaling protein-water interactions for IDPs and multidomain proteins in solution.

### Protein self-association in solution

The observation that multidomain proteins in solution are too compact in Martini 3 simulations suggests that interactions between folded protein domains may be overestimated, at least at the high effective concentration within a single chain. To explore this further, we examined the effect of rescaling protein-protein interactions on the interactions between folded proteins in trans. To this aim, we ran MD simulations of two protein systems that should undergo transient homodimerization, ubiquitin and villin HP36, which we also used in our previous work (***Thomasen et al., 2022***). Ubiquitin self-associates with a *K*_*d*_ of 4.9±0.3 mM based on NMR chemical shift perturbations (***Liu et al., 2012***) and villin HP36 self-associates with a *K*_*d*_ > 1.5 mM based on NMR diffusion measurements (***Brewer et al., 2005***). We ran MD simulations of two copies of the proteins with *λ*_PP_=0.88 and calculated the fraction of the time that the proteins were bound (Fig. 5a). For both proteins *λ*_PP_=0.88 resulted in decreased self-association, and again we found that *λ*_PP_=0.88 gave comparable results to our previously published simulations with *λ*_PW_=1.10 (***Thomasen et al., 2022***). Comparing the simulations with the expected fraction bound based on the experimentally determined *K*_*d*_ values, we found that ubiquitin self-association is likely slightly overestimated with unmodified Martini 3 and slightly underestimated with *λ*_PP_=1.10 and *λ*_PP_=0.88. For villin HP36, all three force fields gave rise to a fraction bound within the expected range. While the overestimated compaction of multidomain proteins suggest that interactions between folded domains may be too strong in Martini 3, our results on the self-association of ubiquitin and villin HP36 do not provide a clear indication that this is the case.

**Figure 5.**
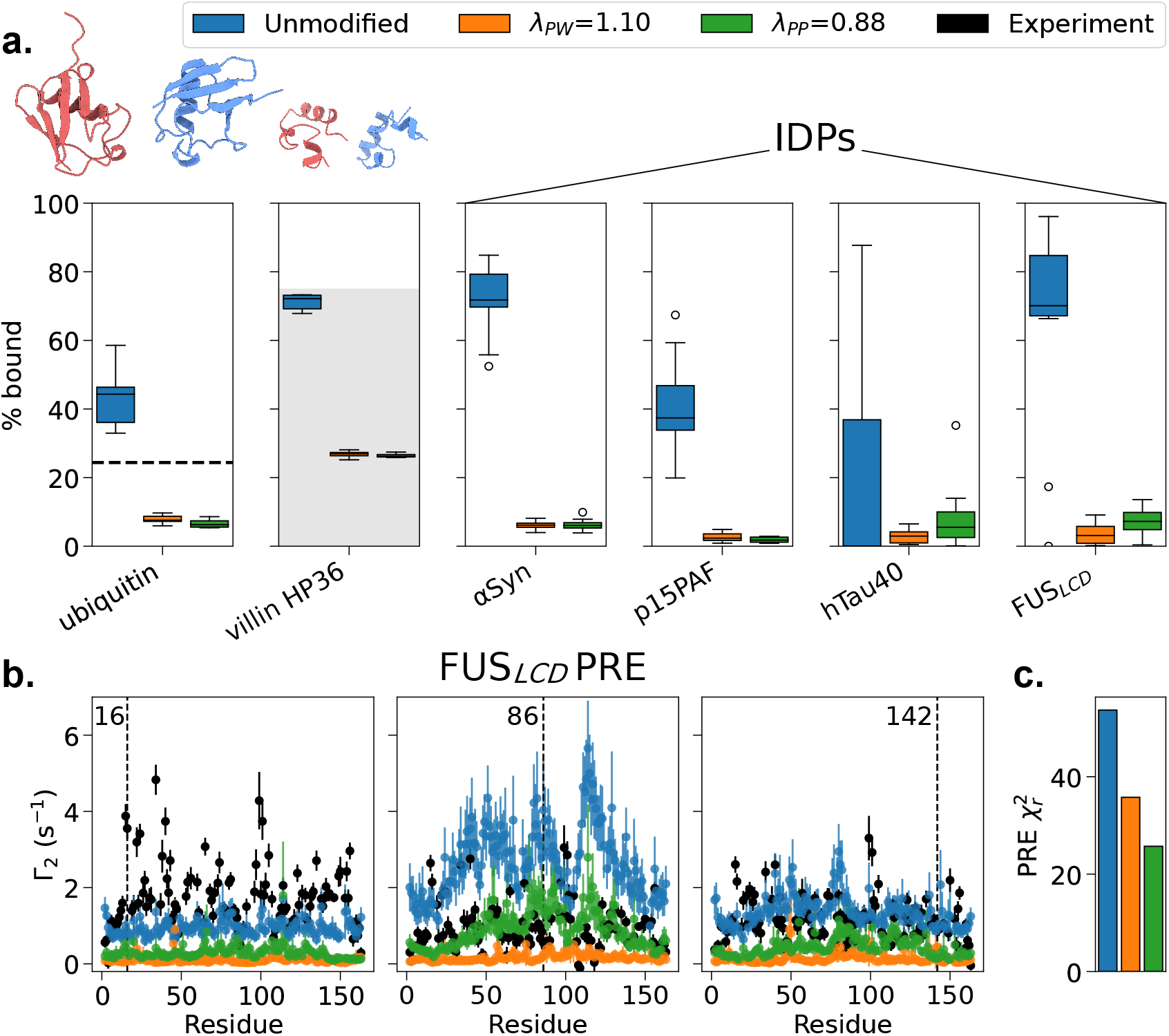
Protein self-association. **a**. Fraction bound calculated from MD simulations of two copies of the folded proteins ubiquitin and villin HP36, and two copies of the IDPs α-synuclein p15PAF, hTau40, and FUS LCD, with unmodified Martini 3 (blue), Martini 3 with protein-water interactions rescaled by *λ*_PW_=1.10 (orange) (taken from ***Thomasen et al. (2022***)), and Martini 3 with protein-protein interactions rescaled by *λ*_PP_=0.88 (green). Box plots show the results of 10 replica simulations. The bound fraction in agreement with Kd = 4.9 mM for ubiquitin self-association is shown as a dashed line (***Liu et al., 2012***). The bound fraction in agreement with a Kd > 1.5 mM for villin HP36 self-association is shown as a shaded gray area (***Brewer et al., 2005***). α-synuclein p15PAF, and hTau40 should not self-associate under the given conditions based on PRE (***Dedmon et al., 2005; Mukrasch et al., 2009***) or SEC-MALLS (***De Biasio et al., 2014***) data, while FUS LCD should transiently self-associate based on PRE data (***Monahan et al., 2017***). Boxplots show the first quartile, median, and third quartile; whiskers extend from the box to the farthest data point lying within 1.5 times the inter-quartile range from the box, and points outside the whiskers are shown individually. **b**. Interchain PREs calculated from the simulations of two copies of FUS LCD from panel a and comparison with experimental PREs (black) (***Monahan et al., 2017***). PREs are shown for the three spin-label sites at residues 16, 86, and 142 marked with dashed black lines. Rotational correlation time *τ*_*c*_ was selected individually for each *λ* to minimize 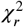. Error bars represent the standard error of the mean. **c**. 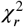 between calculated and experimental PRE data for two copies of FUS LCD shown in panel b.

To further investigate the effect of rescaling protein-protein interactions on protein self-association, we performed simulations of four IDP systems, which we also used in our previous work (***Thomasen et al., 2022***). Specifically, we ran simulations with *λ*_PP_=0.88 of two copies of α-synuclein, hTau40, or p15PAF, which should not self-associate under the given conditions based on PRE (***Dedmon et al., 2005; Mukrasch et al., 2009***) or size-exclusion chromatography-multiangle laser-light scattering (SEC-MALLS) data (***De Biasio et al., 2014***), as well as two copies of the FUS LCD, which should transiently interact under the given conditions based on PRE data (***Monahan et al., 2017***). We then calculated the fraction of time that the proteins were bound in the simulations (Fig. 5a). Again *λ*_PP_=0.88 gave comparable results to our previously published simulations with *λ*_PW_=1.10. The results show that unmodified Martini 3 overestimates the self-association of IDPs, and that both rescaling approaches result in lowered self-association and therefore better agreement with experiments. However, none of the force fields give rise to a clear distinction between the FUS LCD and the three IDPs which should not self-associate, suggesting that Martini 3 does not properly capture specificity in IDP-IDP interactions.

To investigate further how well specific interactions between copies of the FUS LCD were captured, we calculated intermolecular PRE data from our simulations for direct comparison with the experimental PRE data (***Monahan et al., 2017***) (Fig. 5b-c). Simulations with *λ*_PP_=0.88 and *λ*_PW_=1.10 produce similar PREs when calculated with the spin-labels at residue 16 and residue 142, but we observed some discrepancy between the two force fields for PREs calculated with the spin-label at residue 86. These differences could be due to a true difference between the force fields, but may also be due to lack of convergence on the protein-protein contacts, as the bound state is not very populated in the simulations. Both *λ*_PP_=0.88 and *λ*_PW_=1.10 show slight improvement over unmodified Martini 3 based on the 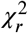 to the experimental PRE data, but none of the force fields fully capture the variation in interactions across the sequence. For example, interactions with the N-terminal region seem to be underestimated with the rescaled force fields based on the PRE data with the spin-label at residue 16, while the interactions with the central region seem to be overestimated with the unmodified force field based on the PRE data with the spin-label at residue 86. The interpretation of the results is complicated by the fact that the rotational correlation time, *τ*_*c*_, providing the best fit to the experimental data is lower for the unmodified force field (1 ns), than for *λ*_PW_=1.10 (8 ns) and *λ*_PP_=0.88 (9 ns), suggesting that the fit of *τ*_*c*_ is absorbing some of the true difference between the force fields. Overall, the comparison with intermolecular PRE data for the FUS LCD is consistent with an improvement in the overall strength of IDP-IDP interactions, but a remaining lack of interaction specificity with the rescaled force fields. The results also show that rescaling protein-protein interactions gives as good or better agreement with the intermolecular PRE data when compared with our previous approach of rescaling protein-water interactions.

### Rescaling protein-water interactions for backbone beads only

While the overall agreement with SAXS experiments was improved for almost all proteins when rescaling protein-protein or protein-water interactions, some proteins were still too expanded or compact with respect to the experimental *R*_*g*_, suggesting that some sequence-specific effects on compaction were not fully captured. We reasoned that sequence-specific effects on the ensemble properties would possibly be better captured if we rescaled only the interactions between the protein backbone and water; this approach could lead to the desired expansion of the proteins while retaining the interactions of the amino acid side chains as originally parameterized. We therefore performed simulations of our set of IDPs and multidomain proteins in which we rescaled ε in the Lennard-Jones potential between all protein backbone and water beads by a factor *λ*_PW-BB_, scanning different values of this parameter, and found *λ*_PW-BB_=1.22 to provide the best agreement with experiments (Fig. S1). However, the simulations of the IDPs and multidomain proteins with *λ*_PW-BB_=1.22 showed similar agreement with experiments as when rescaling all protein-water interactions or protein-protein interactions (Fig. S6-7), and the ensembles resulting from the three rescaling approaches were similar based on *R*_*g*_ distributions (Fig. S2-3) and analyses of the pairwise distances between backbone beads (Fig. S4-5). Given that rescaling of only protein backbone-water interactions did not show any substantial improvement with respect to the previous approaches, and that the strong interactions between the protein backbone and water may have undesirable effects on the behaviour of the hydration shell, we decided not to pursue this approach further.

### Amino acid side chain analogues

We wished to further investigate the symmetry between rescaling of protein-water and protein-protein interactions using simulations of oil/water partitioning, as this was a central approach in the original parameterization of non-bonded interactions in Martini. Inspired by the initial parameterization of Martini proteins, we performed simulations of the cyclohexane/water partitioning of amino acid side chain analogues (***Monticelli et al., 2008***). As rescaling protein-protein interactions should not substantially affect the interactions of amino acids with water or cyclohexane, we ran a single set of simulations to represent both unmodified Martini 3 and *λ*_PP_=0.88, as well as a set of simulations with protein-water interactions rescaled by *λ*_PW_=1.10. We calculated the transfer free energy from cyclohexane to water, Δ*G*_CHEX-W_, from our simulations and compared them with experimentally determined Δ*G*_CHEX-W_-values (Fig. S8) (***Radzicka and Wolfenden, 1988; Monticelli et al., 2008***). The results show that rescaling protein-water interactions by *λ*_PW_=1.10 slightly increases partitioning to the water phase, as would be expected, but the effect is small when compared with the overall discrepancy between simulation and experiment. The two rescaling approaches also provide comparable Pearson correlations with the experimental Δ*G*_CHEX-W_-values (*r*_*Pearson*_ = 0.94±0.03 and *r*_*Pearson*_ = 0.95±0.03 for *λ*_PP_=0.88 and *λ*_PW_=1.10 respectively). We conclude that the results from the oil/water partitioning simulations do not clearly favour one rescaling approach over the other. However, the results illustrate that changes in the non-bonded interactions which have a very modest effect on small molecule partitioning may have a much larger effect on protein-protein interactions and the properties of flexible proteins, highlighting the importance of a direct comparison with experiments that report on protein structure.

Simulations of the dimerization of side chain analogues have previously been used to shed light on similarities and differences across force fields (***de Jong et al., 2012***). We therefore also performed simulations of the self-association of Phe-Phe, Tyr-Phe, Tyr-Tyr, Lys-Asp, and Arg-Asp side chain analogues. Here *λ*_PP_=0.88 and *λ*_PW_=1.10 both result in a small decrease in self-association as measured by the fraction of time bound throughout the simulations (Fig. S9). The two rescaling approaches also give comparable free energy profiles along the center-of-mass (COM) distance, despite the fact that *λ*_PP_=0.88 results in a rebalancing of the Coulomb and Lennard-Jones potentials in the Lys-Asp and Arg-Asp interactions. Comparing with experimentally measured affinities shows that Martini 3 correctly ranks Arg-Asp interactions as stronger than Lys-Asp, and this behaviour is pre-served with both *λ*_PP_=0.88 and *λ*_PW_=1.10 (***Springs and Haake, 1977***). The simulations are also in reasonable agreement with the fraction bound expected from the experimental affinities (***Springs and Haake, 1977***), but perhaps slightly overestimate the strength of the interactions. While the ranking of Tyr-Tyr, Tyr-Phe, and Phe-Phe is consistent with previous analyses of Martini (***de Jong et al., 2012***) and show Phe-Phe to be the strongest in all three versions of Martini 3, analysis of experimental data of disordered proteins suggest that Tyr-Tyr interactions should be stronger than Tyr-Phe or Phe-Phe (***Bremer et al., 2021; Tesei et al., 2021b***). Similarly, measurements of vapour pressure show that benzene-phenol interactions are stronger than benzene-benzene interactions (***Christian and Tucker, 1982***). These results suggest that a rebalancing of aromatic-aromatic interactions in Martini 3 may be necessary to better capture sequence-specific effects in IDPs and multidomain proteins.

### Protein-membrane interactions

In the simulations described above, we found that the effects of increasing protein-water interactions or decreasing protein-protein interactions were very similar. We, however, hypothesized that these two force field modifications could have substantially different effects on systems in which proteins interact with other classes of molecules that are not protein or water. We expected that increased protein-water interactions would result in lower affinity for other molecules, which bind in competition with solvation, while decreased protein-protein interactions would not affect the affinity to the same extent, barring any effects of altering the conformational ensemble.

To examine the effect of rescaling the Lennard-Jones interaction parameters on the affinity of proteins for different biomolecules, we chose to investigate protein interactions with lipid membranes. We had two main motivations for this choice: first, protein-membrane interactions have been thoroughly characterized using Martini (***Yamamoto et al., 2015; Naughton et al., 2016; Srinivasan et al., 2021***); second, Martini has from its early development days in particular been focused on lipid membranes and protein-membrane interactions (***Marrink and Tieleman, 2013; Herzog et al., 2016; Javanainen et al., 2017***).

We therefore performed simulations of peripheral proteins in the presence of lipid bilayers, using both unmodified Martini 3 and the two modified versions, *λ*_PP_=0.88 and *λ*_PW_=1.10, following a protocol we have previously described (***Srinivasan et al., 2021***). In short, we ran unbiased MD simulations starting with the protein at a minimum distance of 3 nm away from the bilayer. Over the course of the MD simulation, the proteins interact, often transiently and reversibly, with the membrane (Fig. S10-11), and membrane binding was quantified as previously described (***Srinivasan et al., 2021***) based on defining bound states when the minimum distance was lower than or equal to 0.7 nm.

To characterize the effect of our rescaling protocol on a broad set of protein-membrane interactions, we selected a diverse set of proteins: (i) one negative control, hen egg-white lysozyme, which is highly soluble in water and is not expected to interact specifically with the membrane in the absence of negatively charged phospholipids (***Howard et al., 1988***); (ii) three peripheral membrane proteins consisting of a single folded domain (Phospholi-pase2, Arf1 in its GTP-bound state, and the C2 domain of Lactadherin) for which we previously characterized the membrane-binding behaviour (***Srinivasan et al., 2021***); (iii) two membrane-binding multidomain proteins: PTEN (1–351), containing a N-terminal Phos-phatase domain and C2 domain that are known to be sufficient for membrane binding, and the Talin FERM domain, that has multiple sub-domains (F0 to F3) and binds to membranes through specific phosphoinositol(4,5)phosphate (PIP2) binding sites present in its F2 and F3 subdomains (***Buhr et al., 2023***); (iv) two intrinsically disordered regions (IDRs) that have been characterized as membrane-binding regions: the N-terminal IDR of TRPV4 (***Goretzki et al., 2023***) and a short C-terminal motif (CTM) of Complexin (***Snead et al., 2014***). For the two IDRs, simulations in solution with both *λ*_PP_=0.88 and *λ*_PW_=1.10 result in expanded ensembles and a larger average value of *R*_*g*_ compared to unmodified Martini 3 (Fig. S12).

As hypothesized, the different force field modifications have different effects on protein-membrane interactions (Fig. 6). In particular, we find that simulations with decreased protein-protein interactions (*λ*_PW_=0.88) provide a similar degree of protein-membrane interaction when compared with unmodified Martini-3. In contrast, simulations with an increased strength of protein-water interactions (*λ*_PW_=1.10) show significantly reduced membrane affinity and binding for all proteins, almost always leading to a complete lack of interactions between the protein and the lipid bilayer. Given that *λ*_PW_=1.10 and *λ*_PP_=0.88 provide a comparably good description of IDPs and multidomain proteins in solution, and that *λ*_PP_=0.88 more accurately retains the specificity and strength of protein-membrane interactions as originally parameterized in Martini 3, we suggest that *λ*_PP_=0.88 is overall a more robust and transferable modification to Martini 3.

**Figure 6.**
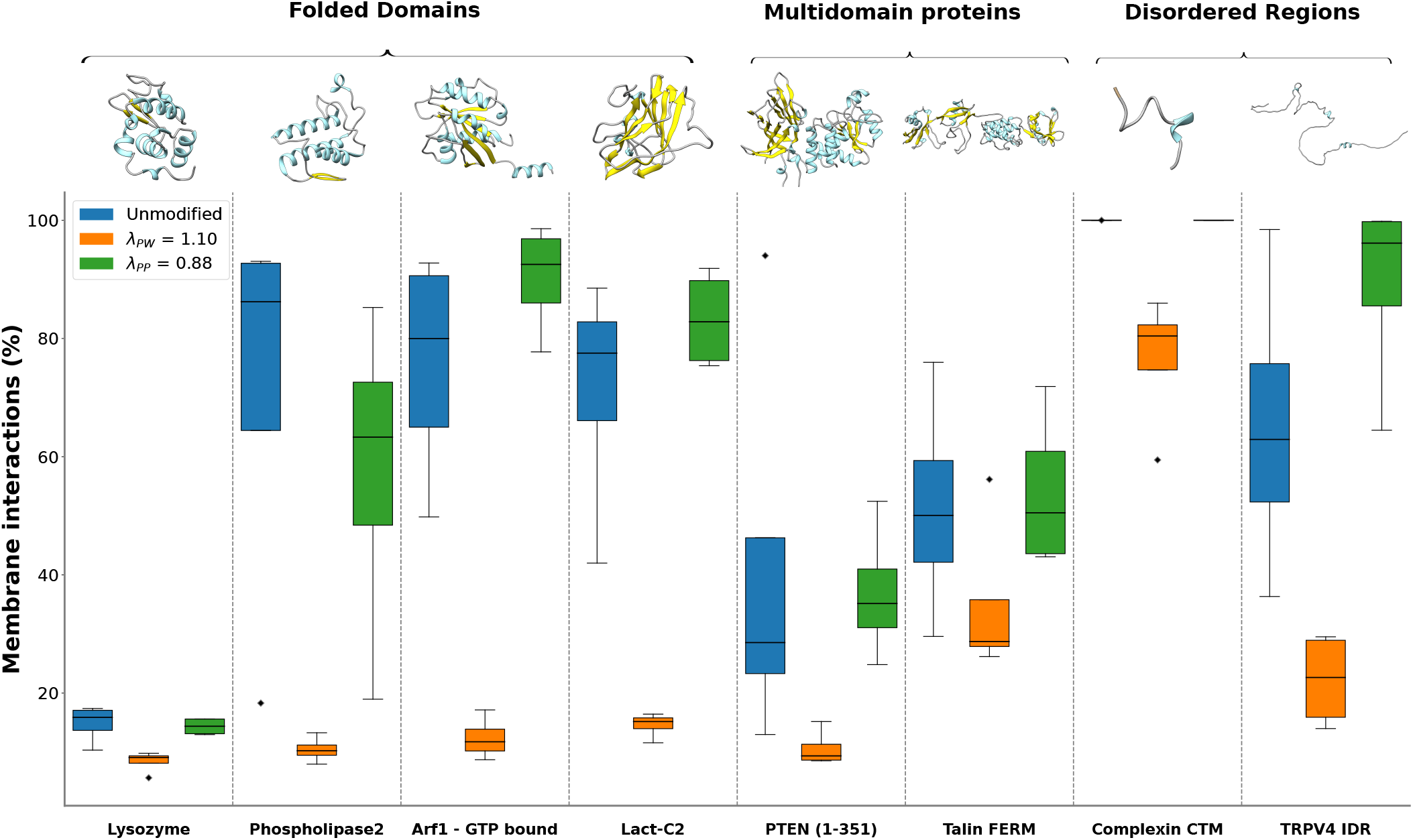
Protein-membrane interactions. MD simulations (four replicas, each 3 µs long) were performed for peripheral membrane proteins, multidomain proteins, and intrinsically disordered regions with appropriate membrane composition (see Methods for details). Simulations were performed with unmodified Martini 3 (blue), protein-water interactions in Martini 3 rescaled by *λ*_PW_=1.10 (orange), and protein-protein interactions in Martini 3 rescaled by *λ*_PP_=0.88 (green). For each system, the corresponding atomistic structure of the protein is shown on top. Boxplots show the first quartile, median, and third quartile; whiskers extend from the box to the farthest data point lying within 1.5 times the inter-quartile range from the box, and points outside the whiskers are shown individually.

### Capturing effects of sequence changes

Having selected *λ*_PP_=0.88 as the preferred force field modification for proteins in solution, we next examined to what extent this force field could capture more subtle sequence effects in IDPs and multidomain proteins.

The *λ*_PP_=0.88 force field provides the same Pearson correlations between experimental and simulation *R*_*g*_ as unmodified Martini 3, initially suggesting that there is no improvement in capturing relative protein-specific differences in *R*_*g*_ (Fig. 3 and 4). To test this for a series of similar proteins with systematic differences in sequence and structure, we selected the mTurq-GS_X_-mNeon proteins, for which the *R*_*g*_ should increase systematically with linker length. We calculated the Pearson correlation between simulation and experimental *R*_*g*_-values for these proteins with the different force fields (Fig. 7a), and found that the simulations with unmodified Martini 3 only provide a small separation of the *R*_*g*_-values as a function of linker length, and therefore give a Pearson correlation coefficient with a high degree of uncertainty based on bootstrapping (*r*_*Pearson*_=0.6±0.6), while the simulations with rescaled interactions allow for a clearer separation of *R*_*g*_ as a function of linker length (*r*_*Pearson*_=0.9±0.1 for both *λ*_PW_=1.10 and *λ*_PP_=0.88). This result suggests that rescaling protein-water or protein-protein interactions allows for a higher sensitivity of ensemble properties to subtle changes in protein sequence and structure, such as differences in interdomain linker length.

**Figure 7.**
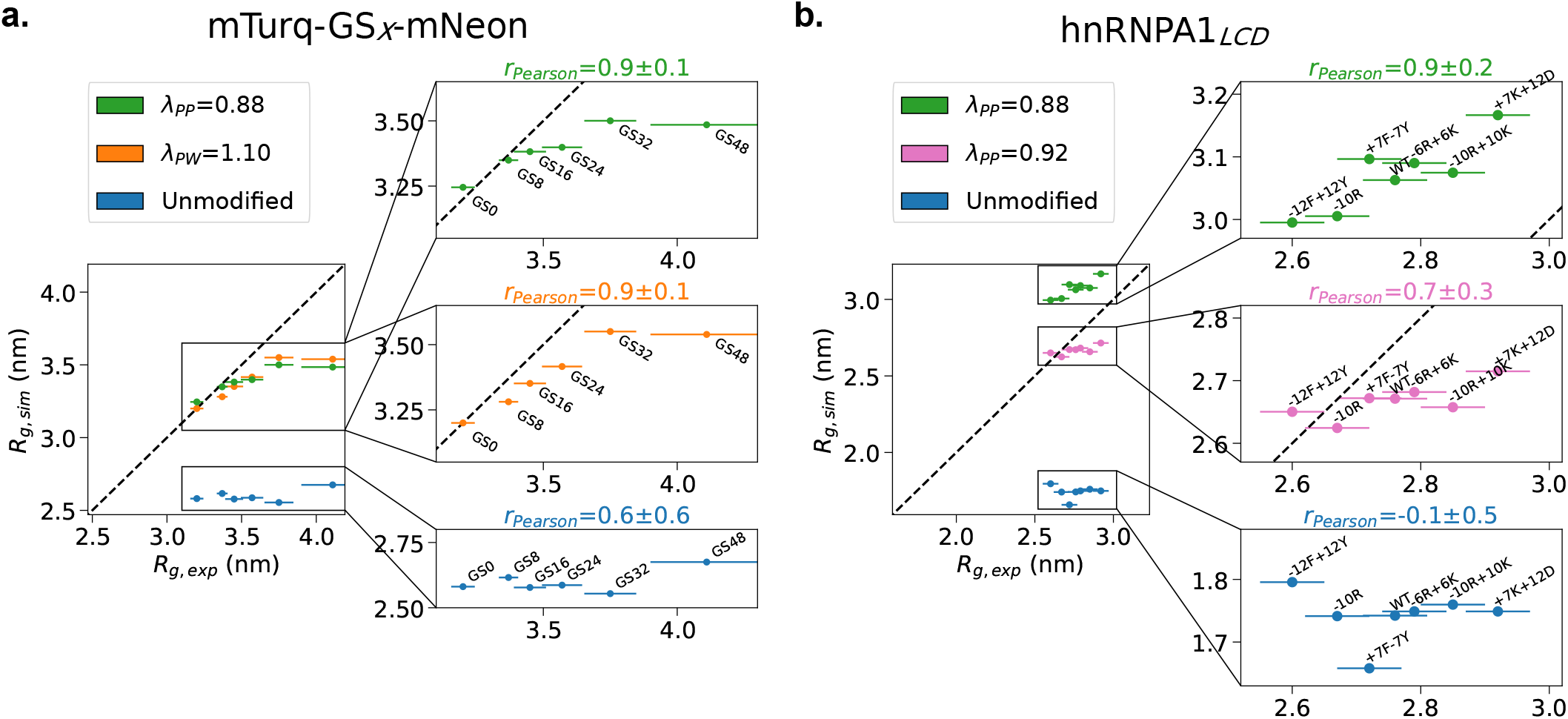
Radii of gyration of mTurq-mNeon and hnRNPA1_LCD_ variants. **a**. *R*_*g*_ calculated from simulations with unmodified Martini 3 (blue), Martini 3 with protein-water interactions rescaled by *λ*_PW_=1.10 (orange), and Martini 3 with protein-protein interactions rescaled by *λ*_PP_=0.88 (green) are plotted against *R*_*g*_ determined by SAXS for mTurquoise2 and mNeonGreen connected by a linker region with the insertion of 0, 8, 16, 24, 32, or 48 GS repeats (***Moses et al., 2024***). **b**. *R*_*g*_ calculated from simulations with unmodified Martini 3 (blue) and Martini 3 with protein-protein interactions rescaled by *λ*_PP_=0.92 (pink) or *λ*_PP_=0.88 (green) are plotted against *R*_*g*_ determined by SAXS for wild-type hnRNPA1_LCD_ and six sequence variants with varied composition of charged and aromatic residues (***Bremer et al., 2022***). We show a zoom-in for each of the force fields along with the given Pearson correlation coefficient with standard error from bootstrapping.

To further investigate the ability of Martini 3 with rescaled protein-protein interactions to capture more subtle sequence effects in IDPs, we performed simulations of six variants of the LCD of hnRNPA1, which have varied composition of charged and aromatic residues while retaining the length of the wild-type sequence (***Bremer et al., 2022***), using unmodified Martini 3 and Martini 3 with *λ*_PP_=0.88. We also performed simulations with *λ*_PP_=0.92, as this provided the optimal agreement with SAXS data for wild-type hnRNPA1. We compared the *R*_*g*_ calculated from the simulations with *R*_*g*_ values measured by SAXS for the six variants and wild-type. As expected based on the results presented above, we found that unmodified Martini 3 substantially underestimates the *R*_*g*_ of all variants (Fig. 7b). While modifying protein-protein interactions by *λ*_PP_=0.88 gives the best results on average across all proteins we studied, it leads to a slight overestimation of the *R*_*g*_ for the wild-type and variants of the LCD from hnRNPA1 (Fig. 7b). If we instead select *λ*_PP_=0.92 as the value of *λ*_PP_ that gives the best result for the wild-type hnRNPA1 LCD (among the values that we examined) we—per construction—find a more accurate level of expansion across the variants. Equally important, we found that unmodified Martini 3 does not accurately capture the variation in *R*_*g*_ associated with the sequence variation (*r*_*Pearson*_=-0.1±0.5), while simulations with *λ*_PP_=0.92 and *λ*_PP_=0.88 result in a more accurate estimate of the effect of the sequence variation on the *R*_*g*_ values (*r*_*Pearson*_=0.7±0.3 and *r*_*Pearson*_=0.9±0.2 respectively). This result suggests that decreasing the strength of protein-protein interactions in Martini 3 improves the sensitivity of IDP ensemble properties to sequence variation.

### Comparison with high-resolution ensembles

Next, we aimed to test the effect of our proposed force field modification by comparing our Martini 3 simulations with ensembles produced by higher resolution models. First, we compared our simulations of α-synuclein with extensive atomistic MD simulations produced with state-of-the-art force fields. We used an ensemble similarity metric based on dimensionality reduction of the pairwise RMSD between ensemble conformers (***Lindorff-Larsen and Ferkinghoff-Borg, 2009; Tiberti et al., 2015***) to quantitatively compare our unmodified and *λ*_PP_=0.88 Martini 3 simulations with atomistic simulations performed with the Amber03ws and Amber99SB-disp force fields (***Robustelli et al., 2018***). We note that both Amber03ws and Amber99SB-disp produce ensembles of α-synuclein which are too compact when compared with *R*_*g*_ from SAXS (***Ahmed et al., 2021***), while the ensemble from our Martini 3 simulation with *λ*_PP_=0.88 is more expanded, in excellent agreement with SAXS. In spite of this discrepancy, the ensemble comparison shows that rescaling protein-protein interactions in Martini 3 by *λ*_PP_=0.88 increases the similarity to the atomistic simulations with both Amber03ws and Amber99SB-disp (Table 1). Interestingly, the *λ*_PP_=0.88 Martini 3 simulation is more similar to both atomistic simulations than the atomistic simulations are to each other, suggesting that the agreement is within the expected variation between force fields.

**Table 1.**
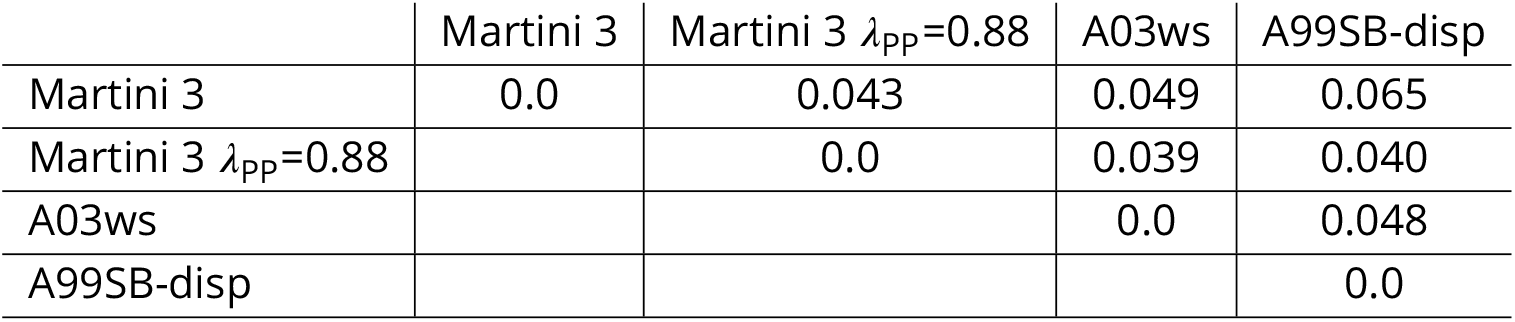
Comparison between Martini and atomistic simulations of α-synuclein. Comparison of unmodified Martini 3 and Martini 3 *λ*_PP_=0.88 simulations of α-synuclein with a 20 µs simulation with the Amber03ws force field and a 73 µs simulation with the Amber99SB-disp force field from ***Robustelli et al. (2018***). The values shown are the Jensen-Shannon divergence calculated with the DRES approach in Encore (***Lindorff-Larsen and Ferkinghoff-Borg, 2009; Tiberti et al., 2015***) based on the Cα RMSD between structures in the two ensembles. The lower bound is 0, corresponding to two identical ensembles, and the upper bound is ln(2) (~0.69).

Next we wished to perform a similar test for a multidomain protein. We used the same approach to quantify the similarity between our unmodified and *λ*_PP_=0.88 Martini 3 simulations of hnRNPA1 with an ensemble that was generated based on data from double electron-electron resonance (DEER) electron paramagnetic resonance, PRE, and SAXS experiments (***Ritsch et al., 2022***). Again, the comparison shows that the Martini 3 simulation with *λ*_PP_=0.88 is more similar to the experimentally derived atomistic ensemble (Table 2). The results from these two test cases suggest that our proposed force field modification of *λ*_PP_=0.88 also improves the agreement with higher resolution simulations and experimentally derived ensemble models.

**Table 2.** Comparison between Martini simulations and an experimentally derived ensemble of hnRNPA1. Comparison of unmodified Martini 3 and Martini 3 *λ*_PP_=0.88 simulations of hnRNPA1 with an atomistic ensemble from ***Ritsch et al. (2022***) generated based on DEER, PRE, and SAXS data (Protein Ensemble Database PED00212). The values shown are the Jensen-Shannon divergence calculated with the DRES approach in Encore (***Lindorff-Larsen and Ferkinghoff-Borg, 2009; Tiberti et al., 2015***) based on the Cα RMSD between structures in the two ensembles. The lower bound is 0, corresponding to two identical ensembles, and the upper bound is ln(2) (~0.69).

| | Martini 3 | Martini 3 $\lambda_{pp}=0.88$ | Expt. ensemble |
| --- | --- | --- | --- |
| Martini 3 | 0.0 | 0.030 | 0.087 |
| Martini 3 $\lambda_{pp}=0.88$ | | 0.0 | 0.077 |
| Expt. ensemble |  |  | 0.0 |

### Protein self-association in the membrane

To test the effect of rescaling protein-protein and protein-water interactions on protein interactions in a lipid membrane environment, we performed simulations of the homodimerization of the transmembrane domain of both EphA1 and ErbB1 from the receptor tyrosine kinase (RTK) domain family, which were used as test systems for Martini 3 (***Souza et al., 2021***). RTKs are a well-studied protein class for protein-protein interactions in a membrane environment and, for both proteins, experimental free energies of association have been determined by Förster resonance energy transfer (FRET) (***Chen et al., 2009; Artemenko et al., 2008***). Our results show that unmodified Martini 3 and Martini 3 with *λ*_*PW*_ =1.10 produce comparable potentials of mean force (PMFs) (Fig. 8), resulting in overestimated Δ*G* of association by ~4 kJ/mol for EphA1 and reasonable agreement with the experimental Δ*G* for ErbB1, consistent with the results from ***Souza et al. (2021***). Rescaling protein-protein interactions by *λ*_*PP*_ =0.88 results in a complete loss of self-association as the PMF profiles becomes repulsive for both proteins (Fig. 8). These results suggest that, while unmodified Martini 3 and Martini 3 with *λ*_*PW*_ =1.10 may slightly overestimate protein-protein interactions in the membrane environment, *λ*_*PP*_ =0.88 results in a substantial underestimation of protein-protein interactions in the membrane, and is likely not a suitable force field modification for studying oligomerization of trans-membrane protein systems.

**Figure 8.**
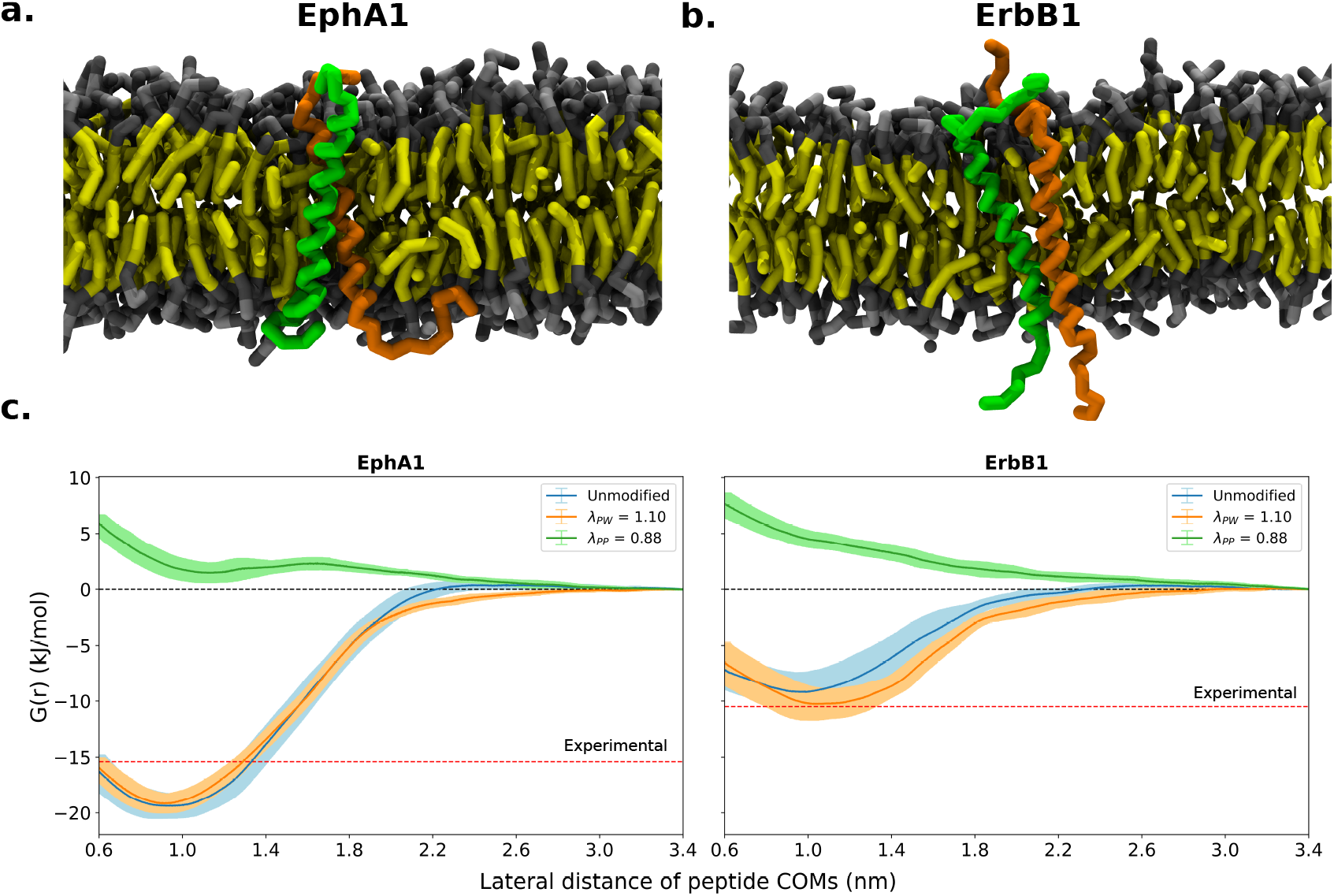
Transmembrane protein self-association. **a-b**. Snapshots of **a**. EphA1 and **b**. ErbB1 simulation systems. The two copies of the protein are shown in green and orange. Lipid heads are shown in gray and lipid tails are shown in yellow. **c**. Potential of mean force profiles for the transmembrane domains of EphA1 and ErbB1 were calculated from simulations with unmodified Martini 3 (blue), Martini 3 with protein-water interactions rescaled by *λ*_PW_=1.10 (orange), and Martini 3 with protein-protein interactions rescaled by *λ*_PP_=0.88 (green). Profiles were aligned to zero at the plateau region at *r*=3.4 nm, indicated by dashed black line. The dashed red line corresponds to experimental values of association free energy (Δ*G*) from FRET experiments for EphA1 and ErbB1. The error represents the standard deviation of four profiles calculated from 2 µs blocks.

## Discussion

We have previously shown that simulations with Martini 3 underestimate the global dimensions of IDPs, and that increasing the strength of protein-water interactions by 10% results in more expanded ensembles and substantially improves the agreement with SAXS data (***Thomasen et al., 2022***). Here, we expanded this approach to a set of 15 multidomain proteins for which SAXS data have been recorded. Our results show that Martini 3 on average provides too compact ensembles of these multidomain proteins, and that, as was the case for IDPs, rescaling protein-water interactions by 10% substantially improves the agreement with SAXS data. We also show that decreasing the strength of interactions between protein beads by 12% results in the same expansion of the ensembles and improved agreement with experiments. We also tested the effect of increasing the strength of interactions between only the protein backbone beads and water, but did not find that this provides any further improvement in the agreement with the experimental data. While the different rescaling approaches provide essentially the same results for proteins in solution, we show that rescaling protein-protein interactions is the preferable option in order to best retain the specificity and strength of protein-membrane interactions as originally parameterized in Martini 3. We note, however, that this change to the force field leads to decreased dimerization of proteins within a membrane environment, and a significant underestimation of free energies of dimerization. Therefore, we suggest that decreasing the strength of protein-protein interactions by 12% is suitable for systems with flexible proteins in solution and in proximity to membranes, but likely not for systems with specific protein-protein interactions in the membrane. An important outcome of our work is also the curation of a set of multidomain proteins with available SAXS data and starting structures for simulations, which can be used for future research in force field assessment and development.

One of the challenges when running Martini 3 simulations of multidomain proteins is selecting which regions to keep folded with the elastic network model and which regions to leave unrestrained. In this work, we manually selected the folded domains in the structures using domain annotations and intuition. It is, however, difficult to know *a priori* whether distinct domains should act as single structural modules due to specific interactions or move freely with respect to one another. Recently, it has been proposed to use the pairwise alignment error output from AlphaFold2 predictions to assign automatically the elastic network restraints (***Jussupow and Kaila, 2023***). In future work, this may provide a more accurate distinction between domains that should be relatively rigid or dynamic with respect to each other. Additionally, replacing the elastic network model with a more flexible structure-based model (***Go, 1983***) may provide the ability to sample both the bound and unbound state in cases where folded domains have specific interactions (***Poma et al., 2017***). In stronger and more specific interdomain interactions, the resolution of Martini 3 may also play a more important role. For example, water-mediated hydrogen-bonding networks would not be captured with the 4-to-1 mapping of water beads. As most of the proteins presented in this work likely do not have very specific interactions between domains, the lack of structured water is presumably not an issue.

Although the simple approach of decreasing the strength of protein-protein interactions uniformly by 12% shows an improvement over unmodified Martini 3 in reproducing the global dimensions of IDPs and multidomain proteins, we note that the agreement with the SAXS data is still not perfect (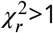 in most cases), and there are systematic out-liers with respect to the experimental *R*_*g*_ values. Although some of the system-specific deviations could potentially be alleviated by e.g. more accurately assigning and modeling the restraints on the folded domains, the overall deviation from the experimental data suggests that a more fundamental rebalancing of non-bonded interactions, and perhaps also CG mapping scheme, is necessary to describe the behaviour of IDPs and multidomain proteins within the Martini framework. Again, we suggest that the data we have collected here will be useful to test any such changes, and the results obtained with *λ*_PP_=0.88 are a useful point of reference for other force field modifications. The increased sensitivity to sequence perturbations observed for the hnNRPA1 sequence variants and the series of mTurq-GS_X_-mNeon proteins also suggests that *λ*_PP_=0.88 could provide a good starting point for rebalancing protein interactions at the amino acid or bead level to improve the specificity in weaker protein-protein interactions.

For other types of systems, it has been suggested that the non-bonded interactions in Martini 3 must be rescaled to a different extent to reach agreement with experimental observations. For example, modifying protein-water interactions in Martini 3 affects the propensity of the disordered LCD of FUS to form condensates in a way that appears to depend on the salt concentration (***Zerze, 2023***), while the insertion of transmembrane helices into the phospholipid bilayer may require decreased protein-water interactions (***Claveras Cabezudo et al., 2023***). Additionally, unmodified Martini 3 has been shown to provide accurate free energies of dimerization for transmembrane proteins (***Souza et al., 2021***). Our results show that this behavior is preserved when rescaling protein-water interactions, whereas decreasing the strength of protein-protein interactions is likely not suitable for systems with specific protein self-association in the membrane. In light of these results, it seems that uniformly rescaling non-bonded interactions may not be able to provide a universally transferable protein model within the Martini framework, and that a more detailed rebalancing of interactions or CG mapping scheme is necessary. Future work could, for example, examine the combined effects of more modest rescaling of protein-protein and protein-water interactions, or focus on secondary-structure dependent force field parameters as recently proposed for another CG force field (***Yamada et al., 2023***).

Overall, however, our results demonstrate that for soluble proteins decreasing the non-bonded interactions between all protein beads by 12% leads to a more accurate balance of interactions while retaining the specificity of protein-membrane interactions. We foresee that our protocol will be a useful starting point to investigate the interactions of IDPs with lipid membranes using chemically transferable MD simulations, and that these investigations will further provide insights into possible strategies on future force field development efforts. Since CG simulations also play an important role in integrative structural biology (***Thomasen and Lindorff-Larsen, 2022***), we also expect that these developments will enable an even tighter link between simulations and experiments to study large and complex biomolecular assemblies.

## Methods

### IDP simulations

We performed MD simulations of a set of 12 IDPs with SAXS data available (Table 3) and five IDPs with intramolceular PRE data available (Table 4) (***Tesei et al., 2021b; Thomasen et al., 2022***) using Gromacs 2020.3 (***Abraham et al., 2015***). We ran simulations with the Martini 3.0 force field (***Souza et al., 2021***) with the well-depth, ε, in the Lennard-Jones potential between all protein beads rescaled by a factor *λ*_PP_ or with ε in the Lennard-Jones potential between all protein backbone and water beads rescaled by a factor *λ*_PW-BB_.

**Table 3.** IDPs with available SAXS data. Number of amino acid residues (*N*_*R*_), experimental *R*_*g*_, temperature (*T*), and salt concentration (*c*_*s*_) used in simulations, and the reference for the SAXS data used.

| Protein | $N_R$ | SAXS $R_g$ (nm) | $T$ (K) | $c_s$ (M) | SAXS ref. |
| --- | --- | --- | --- | --- | --- |
| Hst5 | 24 | $1.34 \pm 0.05$ | 293 | 0.15 | *Jepthah et al. (2019)* |
| Hst5<sub>2</sub> | 48 | $1.77 \pm 0.049$ | 298 | 0.15 | *Fagerberg et al. (2020)* |
| ACTR<sub>AD</sub> | 71 | $2.55 \pm 0.27$ | 278 | 0.2 | *Kjaergaard et al. (2010)* |
| Sic1 | 92 | $2.86 \pm 0.14$ | 293 | 0.2 | *Gomes et al. (2020)* |
| ColN<sub>T</sub> | 98 | $2.82 \pm 0.034$ | 277 | 0.4 | *Johnson et al. (2017)* |
| Tau<sub>K19</sub> | 99 | $3.35 \pm 0.29$ | 288 | 0.15 | *Mylonas et al. (2008)* |
| hnRNPA1<sub>LCD</sub> | 137 | $2.55 \pm 0.1$ | 296 | 0.05 | *Martin et al. (2020)* |
| αSyn | 140 | $3.56 \pm 0.036$ | 293 | 0.2 | *Ahmed et al. (2021)* |
| FhuA | 144 | $3.21 \pm 0.22$ | 298 | 0.15 | *Riback et al. (2017)* |
| Tau<sub>K25</sub> | 185 | $4.06 \pm 0.28$ | 288 | 0.15 | *Mylonas et al. (2008)* |
| CoR<sub>NID</sub> | 271 | $4.72 \pm 0.12$ | 293 | 0.2 | *Cordeiro et al. (2019)* |
| PRN<sub>Nt</sub> | 334 | $4.96 \pm 0.56$ | 298 | 0.15 | *Riback et al. (2017)* |

**Table 4.** IDPs with available PRE data. Number of amino acid residues (*N*_*R*_), temperature (*T*), and salt concentration (*c*_*s*_) used in simulations, and the reference for the PRE data used.

| Protein | $N_R$ | $T$ (K) | $c_s$ (M) | PRE ref. |
| --- | --- | --- | --- | --- |
| αSyn | 140 | 283 | 0.125 | *Dedmon et al. (2005)* |
| hnRNPA2<sub>LCD</sub> | 155 | 298 | 0.005 | *Ryan et al. (2018)* |
| FUS<sub>LCD</sub> | 163 | 298 | 0.15 | *Monahan et al. (2017)* |
| OPN | 220 | 298 | 0.15 | *(Platzer et al., 2011)* |
| hTau40 | 441 | 278 | 0.1 | *(Mukrasch et al., 2009)* |

We generated CG structures using Martinize2 based on initial all-atom structures corresponding to the 95th percentile of the *R*_*g*_-distributions from simulations in ***Tesei et al. (2021b***). Secondary structure and elastic restraints were not assigned for IDPs. Structures were placed in a dodecahedral box using Gromacs editconf and solvated, with NaCl concentrations corresponding to the ionic strength used in SAXS or PRE experiments, using the Insane python script (***Wassenaar et al., 2015***). The systems were equilibrated for 10 ns with a 2 fs time step using the Velocity-Rescaling thermostat (***Bussi et al., 2007***) and Parinello-Rahman barostat (***Parrinello and Rahman, 1981***). Production simulations were run for 40 µs with a 20 fs time step using the Velocity-Rescaling thermostat (***Bussi et al., 2007***) and Parinello-Rahman barostat (***Parrinello and Rahman, 1981***). The simulation temperature was set to match the SAXS or PRE experiment, and the pressure was set to 1 bar. Non-bonded interactions were treated with the Verlet cut-off scheme. A cut-off of 1.1 nm was used for van der Waals interactions. A dielectric constant of 15 and cut-off of 1.1 nm were used for Coulomb interactions. Simulation frames were saved every 1 ns. Molecule breaks from crossing the periodic boundaries were treated with Gromacs trjconv using the flags: -pbc whole -center. Convergence of the simulations was assessed by block-error analysis (***Flyvbjerg and Petersen, 1989***) of *R*_*g*_ calculated from simulation coordinates using the blocking code from: https://github.com/fpesceKU/BLOCKING. All CG trajectories were back-mapped to all-atom structures using a simplified version (***Larsen et al., 2020***) of the Backward algorithm (***Wassenaar et al., 2014***), in which simulation runs are excluded and the two energy minimization runs are shortened to 200 steps.

### Multidomain protein structures

We performed MD simulations of a set of 15 multidomain proteins with SAXS data available (Table 5). We built the initial structure of MyBP-C_MTHB-C2_ based on the NMR structure containing both domains (PDB: 5K6P) (***Michie et al., 2016***). We built the structures of the linear polyubiquitin chains, Ubq_2_, Ubq_3_, and Ubq_4_, based on the crystal structure of the open conformation of Ubq_2_ (PDB: 2W9N) (***Komander et al., 2009***). For Ubq_3_ and Ubq_4_, the linker regions between the original and extended structures were remodelled using Modeller (***Šali and Blundell, 1993***). We built the initial structure of Gal-3 based on the crystal structure of the folded C-terminal domain (PDB: 2NMO) (***Collins et al., 2007***) and the IDR from the AlphaFold structure of full-length Gal3 (AF-P17931-F1) (***Jumper et al., 2021; Tunyasuvunakool et al., 2021***). We built the structure of MyBP-C_C5-C6-C7_ based on the NMR structure of the C5 domain (PDB: 1GXE) (***Idowu et al., 2003***), and the AlphaFold structure of the full-length MyBP-C (AF-Q14896-F1) (***Jumper et al., 2021; Tunyasuvunakool et al., 2021***). We inserted missing residues in the NMR structure of the C5 domain using Modeller (***Šali and Blundell, 1993***). For the mTurq-GS_X_-mNeon constructs, we used structures from Monte-Carlo simulations in ***Moses et al. (2024***) as starting structures for our simulations. To validate the starting structures, we calculated the RMSD between the two fluorescent protein domains and corresponding crystal structures (mTurquoise2 (PDB: 4AR7) (***von Stetten et al., 2012***) and mNeonGreen (PDB: 5LTR) (***Clavel et al., 2016***)) using PyMOL align, which gave an RMSD of 0.2-0.3 Å.

**Table 5.** Multidomain proteins with available SAXS data. Number of amino acid residues (*N*_*R*_), experimental *R*_*g*_, temperature (*T*), and salt concentration (*c*_*s*_) used in simulations, and the reference for the SAXS data used.

| Protein | $N_R$ | SAXS $R_g$ (nm) | $T$ (K) | $c_s$ (M) | SAXS ref. |
| --- | --- | --- | --- | --- | --- |
| MyBP-C<sub>MTHB-C2</sub> | 137 | $1.91 \pm 0.08$ | 277 | 0.15 | *Michie et al. (2016)* |
| Ubq<sub>2</sub> | 162 | $2.2 \pm 0.18$ | 293 | 0.33 | *Jussupow et al. (2020)* |
| Ubq<sub>3</sub> | 228 | $2.62 \pm 0.02$ | 293 | 0.33 | *Jussupow et al. (2020)* |
| Gal-3 | 250 | $2.91 \pm 0.06$ | 303 | 0.04 | *Lin et al. (2017)* |
| TIA1 | 275 | $2.75 \pm 0.05$ | 300 | 0.1 | *Sonntag et al. (2017)* |
| Ubq<sub>4</sub> | 304 | $3.19 \pm 0.09$ | 293 | 0.33 | *Jussupow et al. (2020)* |
| hnRNPA1 | 314 | $3.12 \pm 0.08$ | 300 | 0.15 | *Martin et al. (2021)* |
| MyBP-C<sub>C5-C6-C7</sub> | 328 | $3.75 \pm 0.08$ | 298 | 0.28 | *Nadvi et al. (2016)* |
| hisSUMO-hnRNPA1 | 433 | $3.4 \pm 0.13$ | 300 | 0.1 | *Martin et al. (2021)* |
| mTurq-mNeon | 470 | $3.20 \pm 0.04$ | 293 | 0.15 | *Moses et al. (2024)* |
| mTurq-GS<sub>8</sub>-mNeon | 486 | $3.37 \pm 0.04$ | 293 | 0.15 | *Moses et al. (2024)* |
| mTurq-GS<sub>16</sub>-mNeon | 502 | $3.45 \pm 0.06$ | 293 | 0.15 | *Moses et al. (2024)* |
| mTurq-GS<sub>24</sub>-mNeon | 518 | $3.57 \pm 0.08$ | 293 | 0.15 | *Moses et al. (2024)* |
| mTurq-GS<sub>32</sub>-mNeon | 534 | $3.8 \pm 0.1$ | 293 | 0.15 | *Moses et al. (2024)* |
| mTurq-GS<sub>48</sub>-mNeon | 566 | $4.1 \pm 0.21$ | 293 | 0.15 | *Moses et al. (2024)* |

### Multidomain protein simulations

We ran MD simulations of the set of multidomain proteins using Gromacs 2020.3 (***Abraham et al., 2015***). We ran simulations with the Martini 3.0 force field (***Souza et al., 2021***), as well as several modified versions of Martini 3.0 in which the well-depth, ε, in the Lennard-Jones potential between all protein and water beads was rescaled by a factor *λ*_PW_, ε in the Lennard-Jones potential between all protein beads was rescaled by a factor *λ*_PP_, or ε in the Lennard-Jones potential between all protein backbone and water beads was rescaled by a factor *λ*_PW-BB_. We assigned secondary structure-specific potentials using DSSP (***Kabsch and Sander, 1983***) and Martinize2. The secondary structure of all residues in linkers and IDRs were manually assigned to coil, turn, or bend. We applied an elastic network model using Martinize2 consisting of harmonic potentials with a force constant of 700 kJ mol^-1^ nm^-2^ between all backbone beads within a cut-off distance of 0.9 nm. We removed the elastic network potentials in all linkers and IDRs and between folded domains, so only the structures of individual folded domains were restrained (Table S1). Dihedral and angle potentials between sidechain and backbone beads were assigned using the -scfix flag in Martinize2, but removed in all linkers and IDRs. Structures were placed in a dodecahedral box using Gromacs editconf and solvated, with NaCl concentrations corresponding to the ionic strength used in SAXS experiments, using the Insane python script (***Wassenaar et al., 2015***). The systems were equilibrated for 10 ns with a 2 fs time step using the Berendsen thermostat and Berendsen barostat (***Berendsen et al., 1984***). Production simulations were run for at least 40 µs with a 20 fs time step using the Velocity-Rescaling thermostat (***Bussi et al., 2007***) and Parinello-Rahman baro-stat (***Parrinello and Rahman, 1981***). The simulation temperature was set to match the corresponding SAXS experiment and the pressure was set to 1 bar. Non-bonded interactions were treated with the Verlet cut-off scheme. A cut-off of 1.1 nm was used for van der Waals interactions. A dielectric constant of 15 and cut-off of 1.1 nm were used for Coulomb interactions. Simulation frames were saved every 1 ns. Molecule breaks from crossing the periodic boundaries were treated with Gromacs trjconv using the flags: - pbc whole -center. Convergence of the simulations was assessed by block-error analysis (***Flyvbjerg and Petersen, 1989***) of *R*_*g*_ calculated from simulation coordinates using the blocking code from: https://github.com/fpesceKU/BLOCKING. All CG trajectories were back-mapped to all-atom structures using a simplified version (***Larsen et al., 2020***) of the Backward algorithm (***Wassenaar et al., 2014***), in which simulation runs are excluded and the two energy minimization runs are shortened to 200 steps.

### Simulations of protein self-association in solution

We ran MD simulations of two copies of the two folded proteins ubiquitin and villin HP36, and the four IDPs FUS_LCD_, α-synuclein, hTau40, and p15PAF, as previously described (***Thomasen et al., 2022***), using the Martini 3.0 force field (***Souza et al., 2021***) with the well-depth, ε, in the Lennard-Jones potential between all protein beads rescaled by a factor *λ*_PP_=0.88. We used PDB ID 1UBQ (***Vijay-Kumar et al., 1987***) and PDB ID 1VII (***McKnight et al., 1997***) as starting structures for ubiquitin and villin HP36, respectively. The simulations were set up and run using the same protocol as for IDP simulations. Two copies of ubiquitin, villin HP36, FUS_LCD_, α-synuclein, hTau40, and p15PAF were placed in cubic boxes with side lengths 14.92, 7.31, 40.5, 25.51, 48.02, and 34.15 nm giving protein concentrations of 1000, 8500, 50, 200, 30, and 83.4 µM respectively. NaCl concentrations and temperatures were set according to the corresponding experimental conditions (Table 6 and 7). For ubiquitin and villin HP36 the following steps were also used in the simulation setup: (i) Secondary structure was assigned with DSSP (***Kabsch and Sander, 1983***) in Martinize2. (ii) An elastic network model was applied with Martinize2. The elastic restraints consisted of a harmonic potential of 700 kJ mol^-1^nm^-2^ between backbone beads within a 0.9 nm cut-off. For ubiquitin, we removed elastic restraints from the C-terminus (residues 72–76) to allow for flexibility (***Lindorff-Larsen et al., 2005***). (iv) Dihedral and angular potentials between side chains and backbone beads were added based on the initial structures with the -scfix flag in Martinize2. For ubiquitin, villin HP36, α-synuclein, and p15PAF we ran 10 replica simulations of 40 µs per replica. For hTau40 and FUS_LCD_, we ran 10 replica simulations of 13 µs and 25 µs per replica respectively.

**Table 6.** IDPs with available self-association data. Number of amino acid residues (*N*_*R*_), cubic box side lengths (*d*), simulation temperature (*T*), salt concentration (*c*_*s*_), initial protein concentration (*c*_*p*_) used in simulations, and the reference for the self-association data.

| Protein | $N_R$ | $d$ (nm) | $T$ (K) | $c_s$ (M) | $c_p$ (μM) | Self-association ref. |
| --- | --- | --- | --- | --- | --- | --- |
| αSyn | 140x2 | 25.51 | 283 | 0.125 | 200 | *Dedmon et al. (2005)* |
| FUS<sub>LCD</sub> | 163x2 | 40.5 | 298 | 0.15 | 50 | *Monahan et al. (2017)* |
| p15PAF | 111x2 | 34.15 | 298 | 0.15 | 83.4 | *De Biasio et al. (2014)* |
| hTau40 | 441x2 | 48.02 | 278 | 0.1 | 30 | *Mukrasch et al. (2009)* |

**Table 7.** Folded proteins with available self-association data. Number of amino acid residues (*N*_*R*_), cubic box side lengths (*d*), experimental *K*_*d*_ for self-association, temperature (*T*), salt concentration (*c*_*s*_), initial protein concentration (*c*_*p*_) used in simulations, and the reference for the self-association data.

| Protein | $N_R$ | $d$ (nm) | $K_d$ (mM) | $T$ (K) | $c_s$ (M) | $c_p$ (mM) | Self-association ref. |
| --- | --- | --- | --- | --- | --- | --- | --- |
| Villin HP36 | 36x2 | 7.31 | >1.5 | 298 | 0.15 | 8.5 | *Brewer et al. (2005)* |
| Ubq | 76x2 | 14.92 | $4.9 \pm 0.3$ | 303 | 0.11 | 1.0 | *Liu et al. (2012)* |

We analyzed the population of the bound states in our simulations by calculating the minimum distance between beads in the two protein copies over the trajectory with Gromacs mindist. The fraction bound was defined as the fraction of frames where the minimum distance was below 0.8 nm. For ubiquitin and villin HP36, we calculated the expected fraction of bound protein at the concentrations in our simulations based on the respective *K*_*d*_ -values of 4.9 mM and 1.5 mM determined for self-association (***Liu et al., 2012; Brewer et al., 2005***). The bound fraction was calculated as

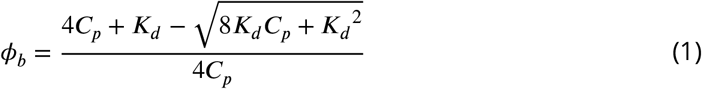

where *ϕ*_*b*_ is the bound fraction, *C*_*p*_ is the concentration of protein in the simulation box (using the average box volume over all simulation trajectories), and *K*_*d*_ is the dissociation constant.

### Amino acid side chain analogue simulations

The Martini 3 parameters for amino acid side chain analogues were produced based on the existing amino acid parameters by simply removing the backbone bead and any potentials or exclusions involving the backbone bead. For simulations of Arg-Asp side chain analogue self-association, the SC1 bead was also removed from Arg (leaving only the SC2 bead of type SQ3p) in order to best emulate the guanidine-acetate system used to measure the experimental affinity (***Springs and Haake, 1977***).

We ran MD simulations of two copies of Tyr and Phe side chain analogues, as well as Tyr-Phe, Arg-Asp, and Lys-Asp side chain analogues using the Martini 3.0 force field (***Souza et al., 2021***) either unmodified or with the well-depth, ε, in the Lennard-Jones potential between all protein and water beads rescaled by a factor *λ*_PW_=1.10 or ε in the Lennard-Jones potential between all protein beads rescaled by a factor *λ*_PP_=0.88. The simulations were set up and run using the same protocol as for IDP simulations. The two side chain analogues were placed in a cubic box with a side length of 5 nm. A NaCl concentration of 150 mM was used for Phe-Phe, Tyr-Phe, Tyr-Tyr simulations. No NaCl was added in the Lys-Asp and Arg-Asp systems. The simulations were run for 100 µs each at 300 K. The fraction bound was calculated from simulations and dissociation constants (*K*_*d*_) using the same approach as for protein self-association in solution described above. Experimental association constants (*K*_*a*_) of 0.4 M^−1^ for Phe-Phe (benzene-benzene) and 0.6 M^−1^ for Phe-Tyr (benzene-phenol) were obtained from ***Christian and Tucker*** (***1982***). Experimental *K*_*a*_-values of 0.31 M^−1^ for Lys-Asp (butylammonium-acetate) and 0.37 M^−1^ for Arg-Asp (guanidine-acetate) were obtained from ***Springs and Haake*** (***1977***). For equation 1, *K*_*d*_ = 1/*K*_*a*_ was used. In order to calculate the free energy profiles along the COM distance, we calculated the COM distance between side chain analogues using Gromacs distance, calculated the probability density using the histogram function in NumPy (***Harris et al., 2020***), and calculated the free energy (in units of *k*_*B*_*T*) as: Δ*G* = −*ln*(*P* (*r*_*COM*_)), where *P* (*r*_*COM*_) is the probability density along the COM distance.

We also ran MD simulations of the cyclohexane/water partitioning of the uncharged amino acid side chain analogues for which experimental transfer free energies were taken from ***Radzicka and Wolfenden*** (***1988***); ***Monticelli et al. (2008***) using unmodified Martini 3.0 and Martini 3.0 with *λ*_PW_=1.10. We prepared a simulation box with 716 copies of Martini 3 cyclohexane (CHEX) and water (W) respectively. For each partitioning simulation, we added a single copy of a side chain analog. Simulations were set up and run using the same protocol as for IDP simulations. Each simulation was run for 100 µs at 300 K.

We calculated the number of contacts between beads in the side chain analogue and CHEX or W over the simulation using Gromacs mindist with a cut-off of 0.8 nm. For a given frame, we considered the side chain analogue as partitioned to the phase with the most contacts (frames with an equal number of CHEX and W contacts were discarded). We then calculated the transfer free energy from cyclohexane to water as:

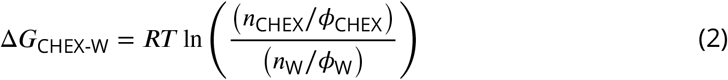

where *R* is the gas constant, *T* is the temperature, *n*_CHEX_ and *n*_W_ are the number of simulation frames where the side chain analogue is in the cyclohexane or water phase respectively, and *ϕ*_CHEX_ and *ϕ*_W_ are the respective volume fractions of the cyclohexane and water phases in the simulations. To determine *ϕ*_CHEX_ and *ϕ*_W_, we calculated the average densities of cyclohexane and water along the z-coordinate in our partitioning simulations of the Ser side chain analogue with unmodified Martini 3 and selected the cut-off between the two phases as the crossover points of the respective densities. The Pearson correlations with experimental transfer free energies were calculated using the pearsonr function in SciPy stats and standard errors were determined with bootstrapping using the bootstrap function in SciPy stats with 9999 resamples (***Virtanen et al., 2020***).

### hnRNPA1 LCD variant simulations

We ran MD simulations of a set of six variants of the hnRNPA1 LCD (−10R, -10R+10K, - 12F+12Y, -6R+6K, +7F-7Y, +7K+12D) for which the *R*_*g*_ has previously been determined by SAXS experiments (***Bremer et al., 2022***). The variants contain substitutions to and from charged and aromatic residues, but have the same sequence length as the wild-type protein, and were selected to have a relatively large deviation in *R*_*g*_ from the wild-type; protein sequences can be found in the supporting information of ***Bremer et al. (2022***). We ran MD simulations with unmodified Martini 3.0 and Martini 3.0 with ε in the Lennard-Jones potential between all protein beads rescaled by a factor *λ*_PP_=0.92 or *λ*_PP_=0.88. Simulations were set up using the same protocol as for the other IDPs described above. The systems were equilibrated for 10 ns with a 2 fs time step using the Berendsen thermostat and Berendsen barostat (***Berendsen et al., 1984***). Production simulations were run for 100 µs with a 20 fs time step using the Velocity-Rescaling thermostat (***Bussi et al., 2007***) and Parinello-Rahman barostat (***Parrinello and Rahman, 1981***). Simulations were run with 150 mM NaCl at 298 K and 1 bar.

### Peripheral membrane protein simulations

We performed MD simulations of one negative control, three peripheral membrane proteins, two multidomain proteins, and two intrinsically disordered regions with lipid bilayers of different compositions (Table 8). We ran simulations with the Martini 3 force field (***Souza et al., 2021***), or with modified force fields in which ε in (i) the Lennard-Jones potential between all protein beads were rescaled by a factor *λ*_PP_=0.88 or (ii) with ε in the Lennard-Jones potential between all protein and water beads rescaled by a factor *λ*_PW_=1.10.

**Table 8.** Membrane-protein systems. Structure (PDB ID), number of amino acid residues (*N*_*R*_) in protein, and lipid composition in the membrane bilayer used in the simulations. Structures starting with AF prefix are AlphaFold-predicted structures (***Varadi et al., 2022***).

| Protein | PDB ID | $N_R$ | Bilayer composition |
| --- | --- | --- | --- |
| Lysozyme | 1AKI | 129 | DOPC |
| Phospholipase2 | 1POA | 118 | DOPC |
| Arf1-GTP bound | 2KSQ | 181 | DOPC |
| Lact-C2 | 3BN6 | 158 | DOPC |
| PTEN (1-351) | AF-F6KD01-F1 | 351 | DOPC:DOPS (8:2) |
| Talin's FERM | 3IVF | 368 | POPC:PIP<sub>2</sub> (10% PIP<sub>2</sub> in upper leaflet) |
| Complexin CTM | Modelled with ColabFold | 16 | POPC:POPS (7:3) |
| TRPV4 IDR | AF-A0A1D5PXA5-F1 | 133 | POPC:DOPS:PIP<sub>2</sub> (7:2:1) |

**Table 9.** Transmembrane proteins with available self-association data. Structure (PDB ID), lipid composition in the membrane bilayer used in simulations, temperature (*T*), salt concentration (*c*_*s*_), and the reference for the self-association data.

| Protein | PDB ID | Bilayer | $T$ (K) | $c_s$ (M) | Self-association ref. |
| --- | --- | --- | --- | --- | --- |
| EphA1 | 2K1L | DLPC | 303 | 0.15 | *Artemenko et al. (2008)* |
| ErbB1 | 2M0B | DLPC | 303 | 0.5 | *Chen et al. (2009)* |

Initial structures of proteins were obtained either from the RCSB database (***Rose et al., 2012***) or from the AlphaFold protein structure database (***Varadi et al., 2022***). For Complexin CTM, we used ColabFold v1.5.2 (***Mirdita et al., 2022***) to model 16-residues long (ATGAFETVKGFFPFGK) disordered region. The N-terminal IDR of TRPV4 (residues 2–134) was taken from the full-length AlphaFold structure of TRPV4 (A0A1D5PXA5). Initial structure of the FERM domains in Talin (PDB:3IVF) had missing residues (134–172), which we modelled using MODELLER (***Webb and Sali, 2016***) via the Chimera interface (***Pettersen et al., 2004***). CG structures of proteins were generated using Martinize2, with DSSP (***Kabsch and Sander, 1983***) flag to assign secondary structures. An elastic network was applied with a harmonic potential of a force constant 700 kJ mol^-1^ nm^-2^ between all backbone beads within a cut-off of 0.8 nm. We removed elastic network potentials between different domains and in linkers and in IDRs of multidomain proteins. Secondary structure and elastic network was not assigned to the two IDRs.

All the lipid bilayers, with initial lateral dimension of 20 nm × 20 nm, were generated using CHARMM-GUI Martini maker (***Qi et al., 2015***), except in the systems where phosphoinositol-(4,5)-phosphate (PIP2) lipids were needed, which instead were generated using the Insane python script (***Wassenaar et al., 2015***). We used the parameter for SAP2_45 lipids (***Borges-Araújo et al., 2021***) to model PIP2 in the bilayer. The bilayers generated from CHARMM-GUI were then minimized and equilibrated following the 6-step equilibration protocol. To compute protein-membrane interactions, systems were generated as previously described (***Srinivasan et al., 2021***), with a minimum distance of 3 nm between any bead of protein and any beads of lipid. Systems were first energy minimized using steepest descent algorithm after which a short MD run of 200 ps was performed with the protein backbone beads restrained. Production simulations (four replicas for each system) were run for 3 µs with a time step of 20 fs using velocity-rescale thermostat (***Bussi et al., 2007***) and Parrinello-Rahman barostat (***Parrinello and Rahman, 1981***).

We performed MD simulation of the two IDRs (Complexin CTM and TRPV4 IDR) in solution with unmodified Martini 3 and both of the modified versions of Martini 3. For these simulations, we took the CG structure and placed it in a cubic box using Gromacs editconf, and solvated and ionized with a concentration of 150 mM of NaCl. Then the system was minimized for 10000 steps with steepest descent algorithm and a short equilibration run was performed with Berendsen thermostat and Berendsen barostat (***Berendsen et al., 1984***) with a time step of 2 fs. Production simulations were run for 10 µs with a 20 fs time-step using Parrinello-Rahman barostat (***Parrinello and Rahman, 1981***) and velocity-rescaling thermostat (***Bussi et al., 2007***). All the simulations were performed with GROMACS 2021.5 (***Abraham et al., 2015***). Initial 100 ns of production run were discarded from all the trajectories for further analysis.

### Simulations of transmembrane protein self-association

We performed simulation of the transmembrane domains of two protein dimers from the RTK family to calculate the free energy of association, Δ*G*, using the Martini 3.0 force field (***Souza et al., 2021***) either unmodified or with the well-depth, ε, in the Lennard-Jones potential between all protein and water beads rescaled by a factor *λ*_PW_=1.10 or ε in the Lennard-Jones potential between all protein beads rescaled by a factor *λ*_PP_=0.88. Simulations were performed with Gromacs 2021.5. PDB 2K1L (***Bocharov et al., 2008***) was used as the starting structure for EphA1. We used Charmm-GUI (***Qi et al., 2015***) to embed the EphA1 dimer in a bilayer of 400 DLPC lipids and 0.5 M NaCl corresponding to the conditions in the reference experiment (***Artemenko et al., 2008***), as in (***Javanainen et al., 2017***). The system was equilibrated using the standard six-step protocol in Charmm-GUI. For ErbB1, the starting structure of the system, based on PDB 2M0B (***Bocharov et al., 2016***), was taken from ***Souza et al. (2021***). The system has 400 DLPC lipids and 0.15 M NaCl corresponding to the conditions in the reference experiment (***Chen et al., 2009***). The system was equilibrated for 50 ns in the NPT ensemble with position restraints of 1000 kJ/(mol nm^2^) on both chain of the dimer.

For both systems, pulling simulations were run for 100 ns at a rate of 0.05 nm/ns. The 2D COM distance between the protein subunits, *r*_*COM*_, was used as the reaction coordinate. Frames ranging from 0.6 nm to 3.4 nm with a spacing of 0.2 nm were extracted from the pulling simulation trajectories as umbrella sampling windows, to be consistent with previous work (***Souza et al., 2021***). A spring constant of 400 kJ/(mol nm^2^) was applied as the umbrella potential in production runs. The temperature was maintained at 303 K separately for peptides, lipids, and solvents and semi-isotropic pressure coupling was applied at 1 bar. The production run was performed for 10 µs in each window using a time-step of 20 fs and the Gromacs wham tool (***Hub et al., 2010***) was used to obtain the PMF. The error in PMF plots represents the standard deviation of 4 profiles calculated from 2 µs blocks, with the first 2 µs were discarded. All PMFs plateaued before *r*_*COM*_ =3.4 nm and were aligned to zero at this value. Δ*G* values were estimated from the minimum of zero aligned PMFs for comparison with experimental values.

### SAXS calculations

We extracted 20,000 evenly distributed frames from each back-mapped trajectory to calculate SAXS profiles using Pepsi-SAXS (***Grudinin et al., 2017***). To avoid overfitting the parameters for the contrast of the hydration layer (*δρ*) and the displaced solvent (*r*0) by fitting them individually to each structure, we used the fixed values for these parameters determined in ***Pesce and Lindorff-Larsen*** (***2021***). We globally fitted the scale and constant background with least-squares regression weighted by the experimental errors using Scikit-learn (***Pedregosa et al., 2011***). To assess the agreement between the experimental SAXS profiles and those calculated from simulations, we calculated the 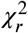 between the ensemble-averaged calculated SAXS intensities (*I*_*eale*_) and the experimental SAXS intensity (*I*_*exp*_):

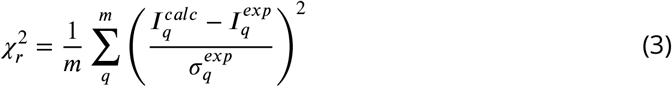

where *(σ*^*exp*^ is the error of the experimental SAXS intensity and *m* is the number of measured SAXS intensities. We used the Bayesian Indirect Fourier Transform algorithm (BIFT) to rescale the errors of the experimental SAXS intensities, in order to obtain a more consistent error estimate across the different proteins (***Hansen, 2000; Larsen and Pedersen, 2021***).

### PRE calulcations

We used the DEER-PREdict software (***Tesei et al., 2021a***) to calculate intrachain PREs from the back-mapped trajectories of α-synuclein, FUS_LCD_,hnRNPA2_LCD_, OPN and hTau40 (Table 4), and interchain PREs from the back-mapped trajectories of two copies of FUS_LCD_. DEER-PREDICT uses a rotamer library approach to model the MTSL spin-label (***Polyhach et al., 2011***) and a model-free formalism to calculate the spectral density (***Iwahara et al., 2004***). We assumed an effective correlation time of the spin label, *τ* _*t*_, of 100 ps, a molecular correlation time, *τ* _*e*_, of 4 ns (***Gillespie and Shortle, 1997***), a transverse relaxation rate for the diamagnetic protein of 10 *s*^−1^ and a total INEPT time of the HSQC measurement of 10 ms (***Battiste and Wagner, 2000***). For the simulations of two copies of FUS_LCD_, *τ* _*e*_ was not fixed to 4 ns. We instead scanned values of *τ* _*e*_ from 1 to 20 ns in steps of 1 ns and selected the *τ* _*e*_ that minimized the 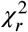 to the experimental PRE data for each force field. The optimal values were 1 ns, 8 ns, and 9 ns for unmodified Martini 3, *λ* _PW_=1.10 and *λ* _PP_=0.88 respectively. The agreement between calculated and experimental PREs was assessed by calculating the *χ* ^2^ over all spin-label positions,

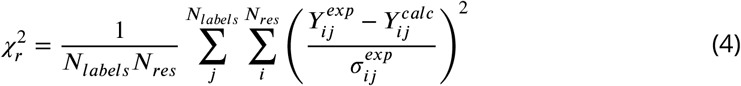

where *N*_*labels*_ and *N*_*res*_ are the number of spin-labels and residues, 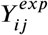 and 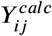 are the experimental and calculated PRE rates for label *J* and residue *i*, and 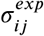 is the experimental error of the PRE rate for label *J* and residue *i*. For the simulations of two copies of the FUS_LCD_, the 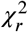 was calculated as an average over the 10 replica simulations.

### Radii of gyration

We calculated the *R*_*g*_ from CG simulation trajectories using Gromacs gyrate (***Abraham et al., 2015***) and calculated the error of the average *R*_*g*_ using block-error analysis (***Fly-vbjerg and Petersen, 1989***) (https://github.com/fpesceKU/BLOCKING). Experimental *R*_*g*_-values and corresponding error bars were calculated from SAXS profiles by Guinier analysis using ATSAS AUTORG with default settings (***Petoukhov et al., 2007***), except in the case of the hnRNPA1_LCD_ variants, for which we used the *R*_*g*_-values reported in ***Bremer et al. (2022***), which were determined from SAXS data using an empirical molecular form factor approach. Pearson correlation coefficients were calculated using the pearsonr function in SciPy stats and standard errors were determined with bootstrapping using the boot-strap function in SciPy stats with 9999 resamples (***Virtanen et al., 2020***).

### Principal component analysis

We used PCA based on the pairwise distances between backbone beads to compare our unmodified, *λ* _PW_=1.10, *λ* _PP_=0.88, and *λ* _PW-BB_=1.22 Martini 3 simulations of IDPs and multidomain proteins. PCA was performed with PyEMMA (***Scherer et al., 2015***). For each protein, all four ensembles were pooled for PCA in order to project into the same two principal components. For all IDPs except PRN_NT_ and CoR_NID_, and for the multidomain proteins Gal-3, MyBP-C_MTHB-C2_, Ubq_2_ and Ubq_3_, the pairwise distances between all back-bone beads were used as features for PCA. For the remaining proteins, the pairwise distances between every 10th backbone bead were used as features for PCA.

### Comparison with atomistic ensembles

We compared our unmodified Martini 3 and *λ* _PP_=0.88 Martini 3 simulations of α-synuclein with a 20 µs simulation with the Amber03ws force field and a 73 µs simulation with the Amber99SB-disp force field from ***Robustelli et al. (2018***). Because of problems caused by interactions between periodic images of the protein in the originally published Amber99SB-disp simulation, we used the corrected version of the simulation also used in ***Ahmed et al. (2021***). We also compared our unmodified Martini 3 and *λ* _PP_=0.88 Martini 3 simulations of hnRNPA1 with an atomistic ensemble from ***Ritsch et al. (2022***) (Protein Ensemble Database PED00212). We used the dimensionality reduction ensemble similarity (DRES) approach in Encore (***Lindorff-Larsen and Ferkinghoff-Borg, 2009; Tiberti et al., 2015***) implemented in MDAnalysis (***Michaud-Agrawal et al., 2011***) to quantify the ensemble similarity based on Cα RMSD. We used the all-atom back-mapped versions of our Martini simulations (described above). For hnRNPA1, we used every 10th frame from our Martini simulations for a total of 4001 frames per simulation and all structures from the atomistic ensemble. Different constructs of hnRNPA1 were used for our simulations and the ***Ritsch et al. (2022***) ensemble, so the Encore DRES calculations were only performed for residues 2-258, which are identical in both constructs (***Martin et al., 2021; Ritsch et al., 2022***). For α-synuclein, we used every 10th frame from each simulation for a total of 4001 frames per Martini simulation, 2998 frames from the Amber03ws simulation, and 2998 frames from the Amber99sb-disp simulation.

## Data availability

The data generated for this paper is available via https://github.com/KULL-Centre/_2023_Thomasen_Martini. Simulation data and starting structures for simulations are available at https://doi.org/10.5281/zenodo.8010043. Data for protein membrane simulations are available at https://zenodo.org/record/8154919. Force field files for Martini 3 with interactions between protein beads rescaled by *λ*_PP_=0.88 are available at https://github.com/KULL-Centre/_2023_Thomasen_Martini/tree/main/force_field

## Code availability

Code and scripts used for this paper is available via https://github.com/KULL-Centre/_2023_Thomasen_Martini.

## Acknowledgments

We acknowledge the use of computational resources from Computerome 2.0, the ROBUST Resource for Biomolecular Simulations (supported by the Novo Nordisk Foundation grant no. NF18OC0032608), and the core facility for biocomputing at the Department of Biology. This research was supported by the Lundbeck Foundation BRAINSTRUC initiative (R155-2015-2666 to K.L.-L.) and the PRISM (Protein Interactions and Stability in Medicine and Genomics) centre funded by the Novo Nordisk Foundation (NNF18OC0033950, to K.L.-L.). SV and AK acknowledge support by the Swiss National Science Foundation through the National Center of Competence in Research Bio-Inspired Materials. This work was supported by grants from the Swiss National Supercomputing Centre (CSCS) under project ID s1176 and s1251.

## Author contributions

F.E.T., S.V. and K.L.-L. conceived the overall study. F.E.T and T.S. performed and analysed simulations of proteins in water under the supervision of K.L.-L., and A.K. and S.S. performed and analysed simulations of proteins interacting with membranes under the supervision of S.V.. F.E.T. wrote the first draft of the manuscript with input from K.L.-L. All authors contributed to the writing of the manuscript.

