## Supporting Figures and Table for "Rescaling protein-protein interactions improves Martini 3 for flexible proteins in solution"

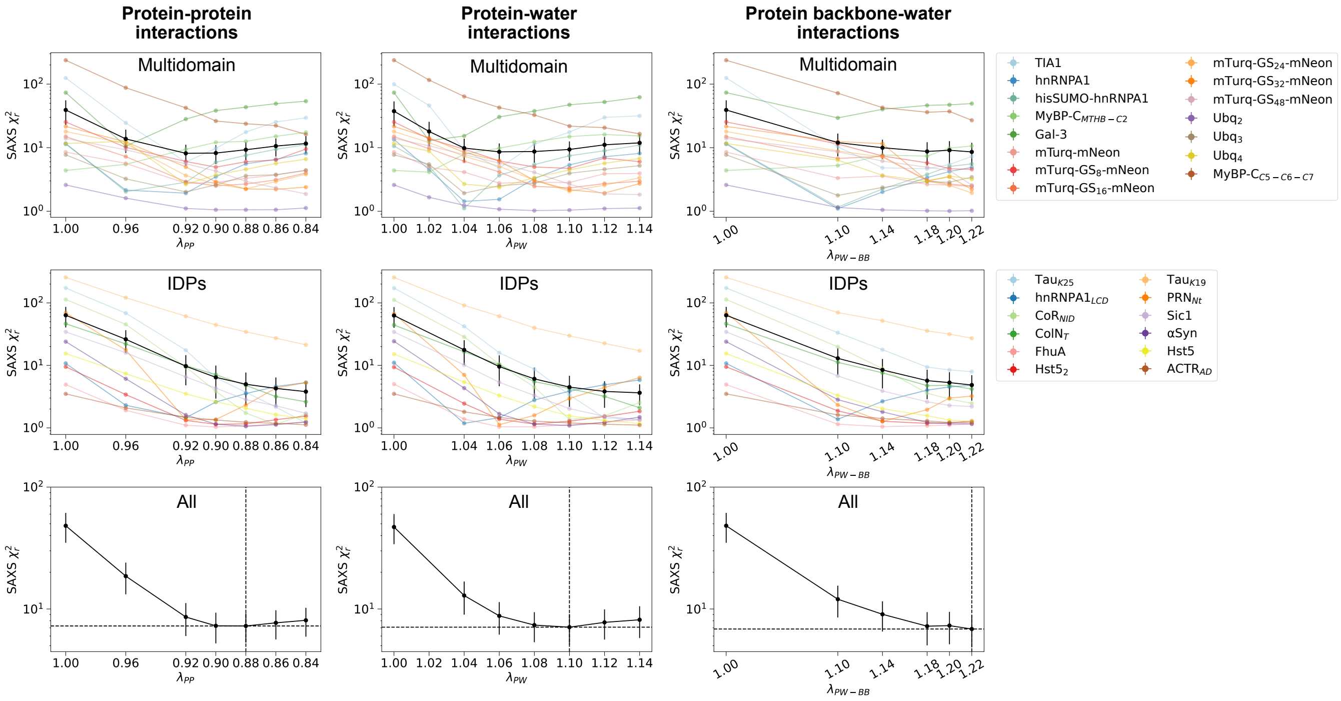


**Figure S1. Scan of rescaling factor $\lambda$ for different sets of non-bonded interactions.** Reduced $\chi^2$ between experimental SAXS intensities and SAXS intensities calculated from Martini 3 simulations in which (left) the well-depth, $\epsilon$, in the Lennard-Jones potential between all protein beads was rescaled by a factor $\lambda_{\mathrm{PP}}$, (middle) $\epsilon$ in the Lennard-Jones potential between all protein and water beads was rescaled by a factor $\lambda_{\mathrm{PW}}$, or (right) $\epsilon$ in the Lennard-Jones potential between all protein backbone and water beads was rescaled by a factor $\lambda_{\mathrm{PW\text{-}BB}}$. The scan of $\lambda$-values is shown for a set of 15 multidomain proteins (top), a set of 12 IDPs (middle), and averaged over all 27 multidomain proteins and IDPs (bottom). Mean and standard error of the mean over different proteins are shown in black. The selected value of $\lambda$, which gives the minimum average $\chi_r^2$ to the SAXS data over all proteins, is indicated with dashed lines in the bottom plot. Note the logarithmic scale. Protein-water rescaling simulations of IDPs and TIA1, hnRNPA1, and hisSUMO-hnRNPA1 are taken from ***Thomasen et al.*** (***2022***).

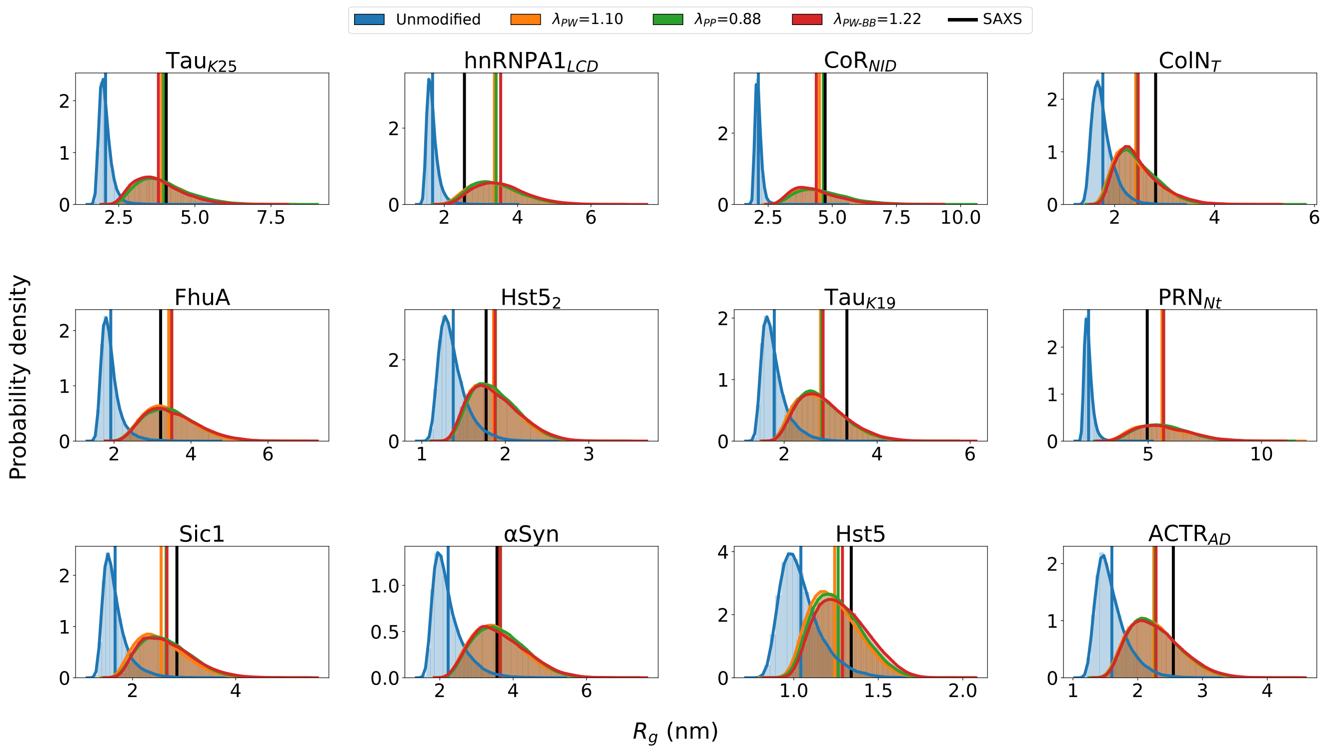


**Figure S2.** **$R_g$-distributions from simulations of IDPs.** $R_g$-distributions for 12 IDPs calculated from simulations with unmodified Martini 3 (blue) or Martini 3 with interactions between protein and water beads rescaled by $\lambda_{\mathrm{PW}}$=1.10 (orange, simulations from ***Thomasen et al.*** (***2022***)), between protein beads rescaled by $\lambda_{\mathrm{PP}}$=0.88 (green), or between protein backbone and water beads rescaled by $\lambda_{\mathrm{PW\text{-}BB}}$ (red). Average values are shown as vertical lines. The experimental $R_g$ from Guinier analysis of SAXS data is shown in black.

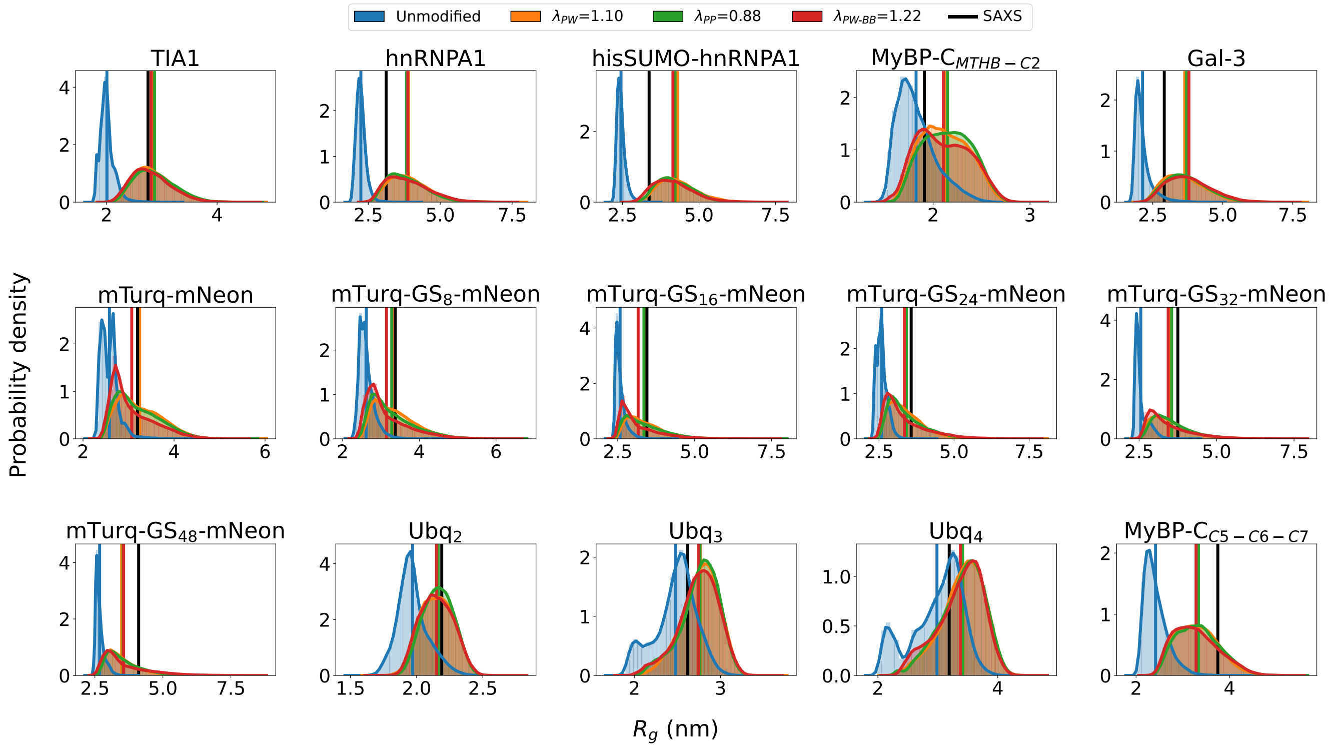


**Figure S3.** **$R_g$-distributions from simulations of multidomain proteins.** $R_g$-distributions for 15 multidomain proteins calculated from simulations with unmodified Martini 3 (blue) or Martini 3 with interactions between protein and water beads rescaled by $\lambda_{\text{PW}}$=1.10 (orange, simulations from ***Thomasen et al.*** (***2022***) for TIA1, hnRNPA1, and hisSUMO-hnRNPA1), between protein beads rescaled by $\lambda_{\text{PP}}$=0.88 (green), or between protein backbone and water beads rescaled by $\lambda_{\text{PW-BB}}$ (red). Average values are shown as vertical lines. The experimental $R_g$ from Guinier analysis of SAXS data is shown in black.

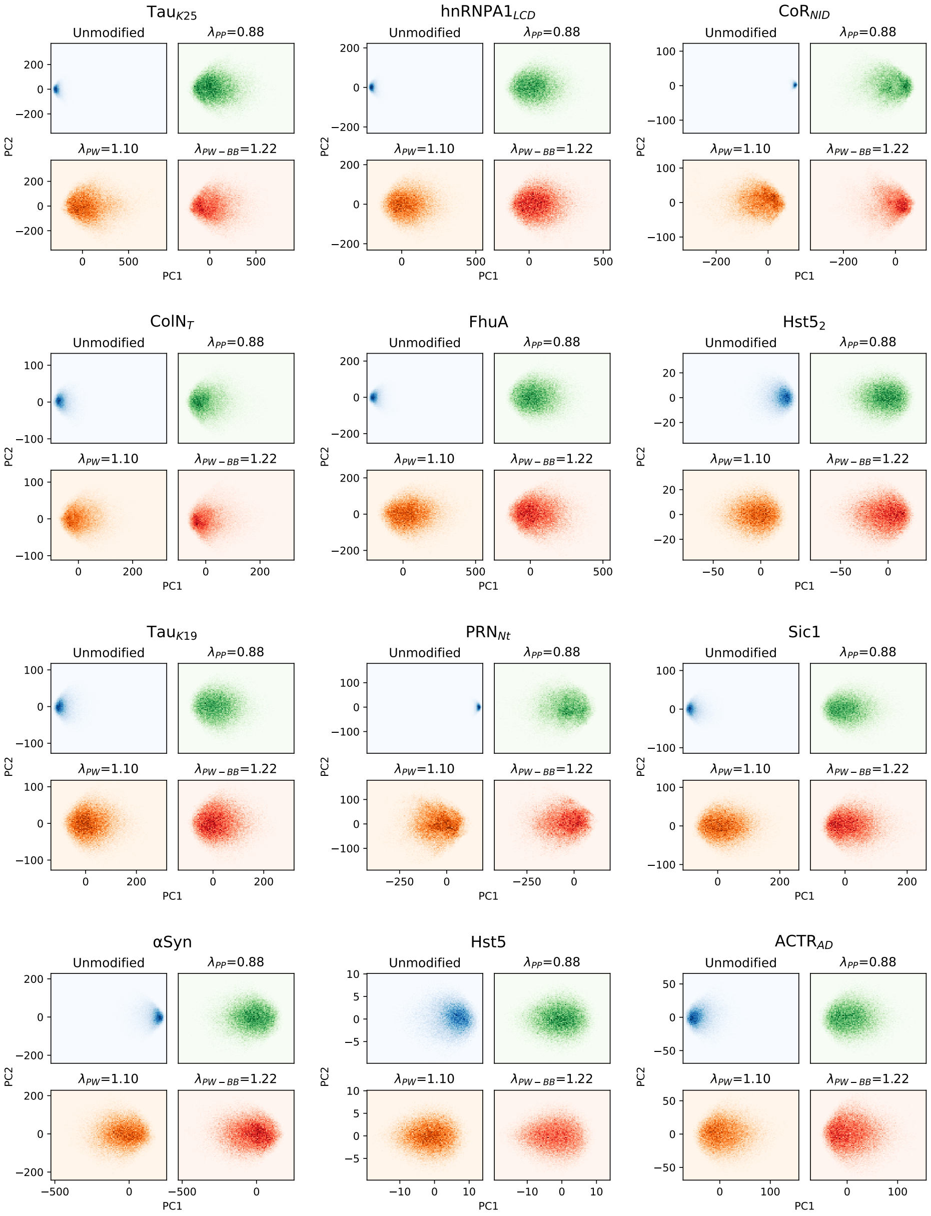


**Figure S4. Principal component analysis of pairwise backbone bead distances for IDPs.** Principal component analysis (PCA) of pairwise C$\alpha$ distances calculated from simulations of a set of 12 IDPs with unmodified Martini 3 (blue) or Martini 3 with interactions between protein and water beads rescaled by $\lambda_{\text{PW}}$=1.10 (orange, simulations from ***Thomasen et al.*** (***2022***)), between protein beads rescaled by $\lambda_{\text{PP}}$=0.88 (green), or between protein backbone and water beads rescaled by $\lambda_{\text{PW-BB}}$ (red). For each protein, simulations with the four different force fields were pooled for PCA to allow for projection onto the same two principal components.

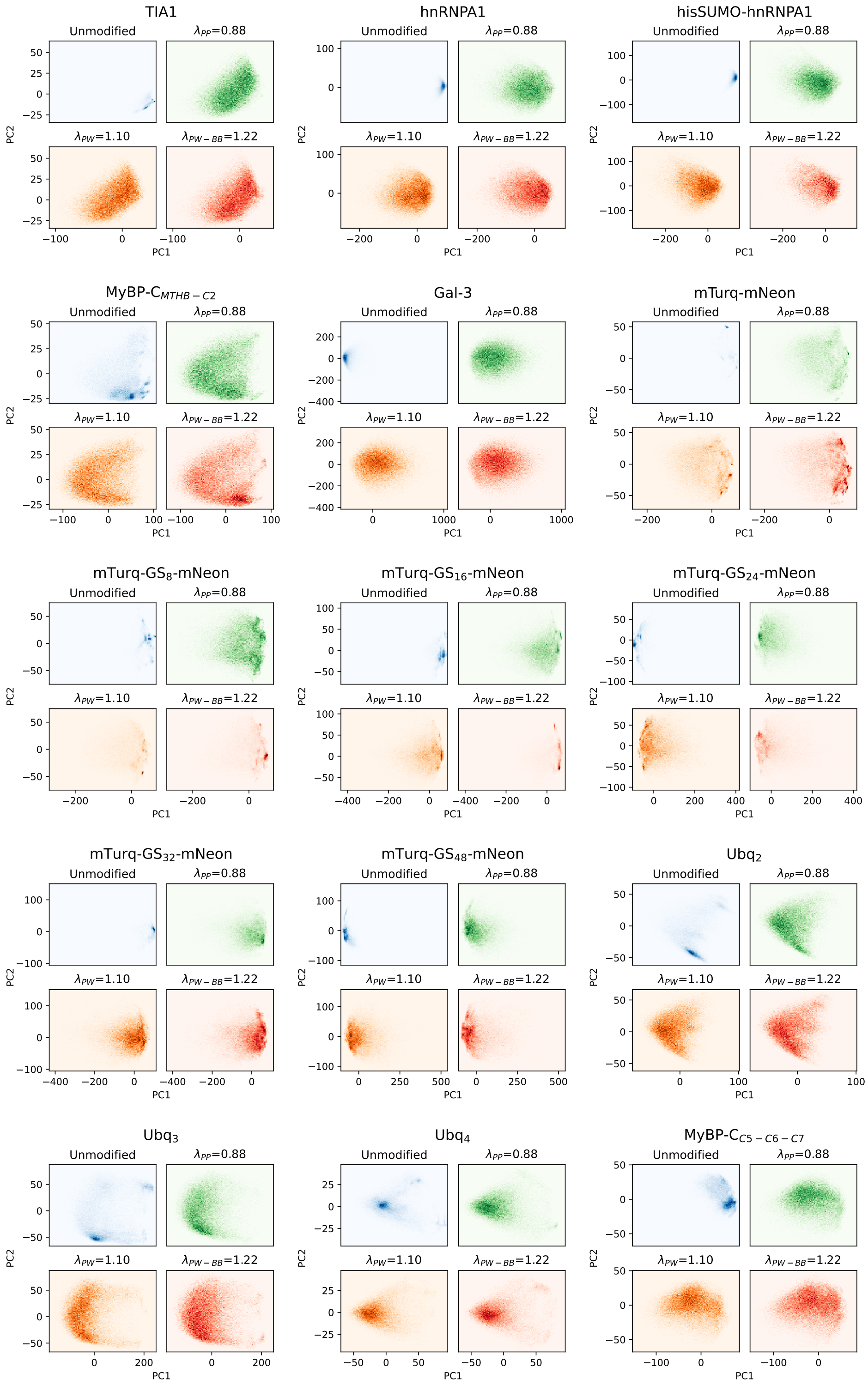


**Figure S5. Principal component analysis of pairwise backbone bead distances for multidomain proteins.** PCA of pairwise backbone bead distances calculated from simulations of a set of 15 multidomain proteins with unmodified Martini 3 (blue) or Martini 3 with interactions between protein and water beads rescaled by $\lambda_{\mathrm{PW}}$=1.10 (orange, simulations from ***Thomasen et al.*** (***2022***) for TIA1, hnRNPA1, and hisSUMO-hnRNPA1), between protein beads rescaled by $\lambda_{\mathrm{PP}}$=0.88 (green), or between protein backbone and water beads rescaled by $\lambda_{\mathrm{PW\text{-}BB}}$ (red). For each protein, simulations with the four different force fields were pooled for PCA to allow for projection onto the same two principal components.

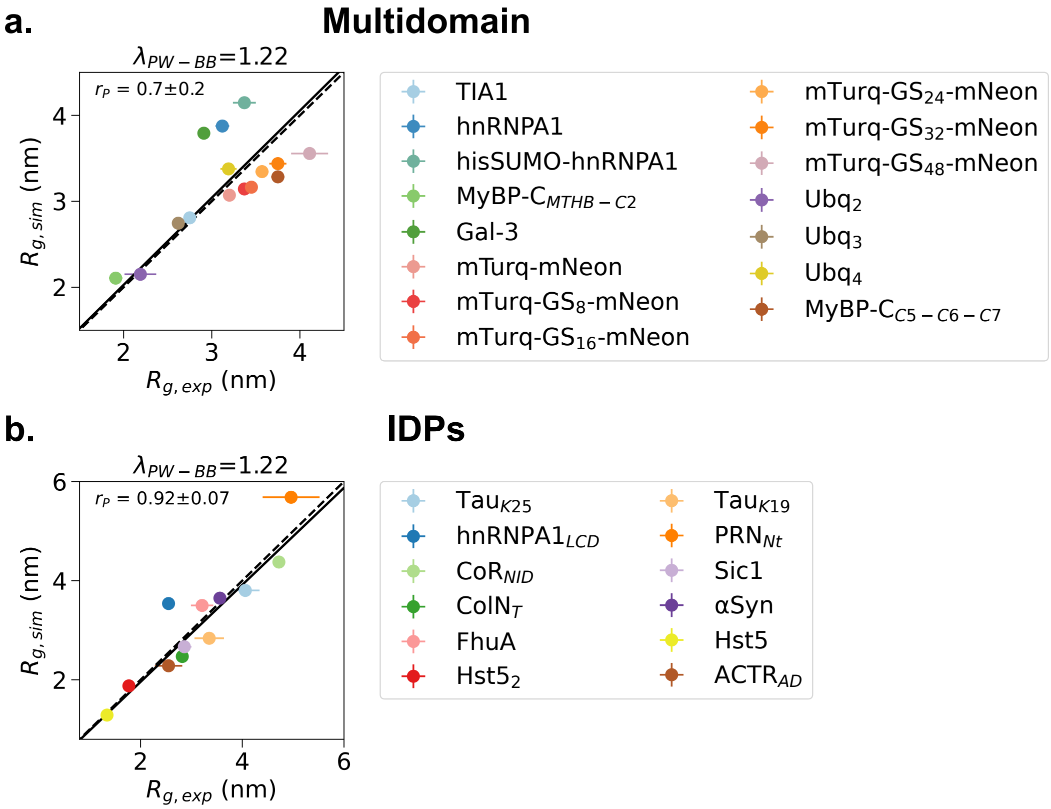


**Figure S6. Agreement between simulation and experimental $R_g$ with rescaled protein backbone-water interactions. a.** $R_g$ calculated from Martini 3 simulations with protein backbone-water interactions rescaled by $\lambda_{\text{PW-BB}}$=1.22 plotted against $R_g$ determined from Guinier fit for 15 multidomain proteins. **c.** $R_g$ calculated from Martini 3 simulations with protein backbone-water interactions rescaled by $\lambda_{\text{PW-BB}}$=1.22 plotted against $R_g$ determined from Guinier fit for 12 IDPs. The diagonal is shown as a dashed line and a linear fit with intercept 0 weighted by experimental errors is shown as a solid line. Pearson correlation coefficients with standard errors from bootstrapping are shown on the plots.

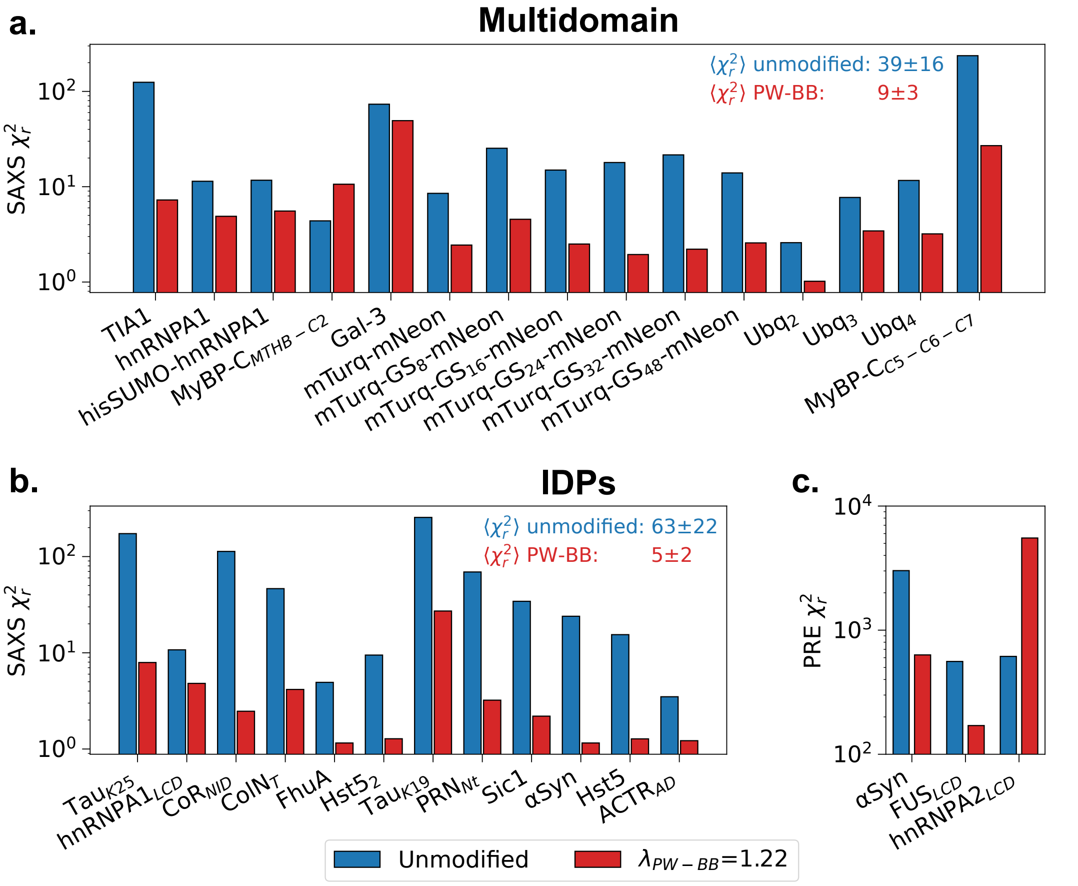


**Figure S7. Agreement between simulations and SAXS or PRE data with rescaled protein backbone-water interactions. a.** Reduced $\chi^2$ between experimental SAXS intensities and SAXS intensities calculated from unmodified Martini 3 simulations (blue) and Martini 3 simulations with protein backbone-water interactions rescaled by $\lambda_{\text{PW-BB}}$=1.22 (red) for at set of 15 multidomain proteins. Mean and standard error of the mean over all proteins are shown on the plot. Note the logarithmic scale. **b.** Reduced $\chi^2$ between experimental SAXS intensities and SAXS intensities calculated from unmodified Martini 3 simulations (blue) and Martini 3 simulations with protein backbone-water interactions rescaled by $\lambda_{\text{PW-BB}}$=1.22 (red) for at set of 12 IDPs. Mean and standard error of the mean over all proteins are shown on the plot. Note the logarithmic scale. **c.** Reduced $\chi^2$ between experimental PRE data and PRE data calculated from unmodified Martini 3 simulations (blue) and Martini 3 simulations with protein backbone-water interactions rescaled by $\lambda_{\text{PW-BB}}$=1.22 (red).

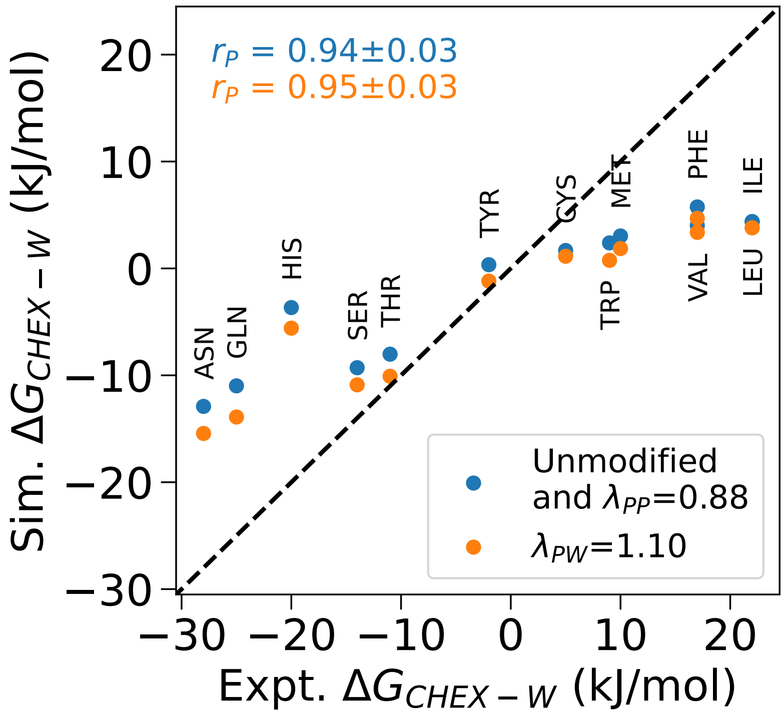


**Figure S8. Cyclohexane/water partitioning of amino acid side chain analogs.** Transfer free energies from cyclohexane to water ($\Delta G_{CHEX-W}$) calculated from simulations of the cyclohexane/water partitioning of amino acid side chain analogs with unmodified Martini 3 (same as with protein-protein interactions rescaled by $\lambda_{PP}$=0.88, blue) and protein-water interactions in Martini 3 rescaled by $\lambda_{PW}$=1.10 (orange) plotted against experimental values of $\Delta G_{CHEX-W}$ (***Radzicka and Wolfenden, 1988***; ***Monticelli et al., 2008***). Pearson correlation coefficients with standard errors from bootstrapping are shown on the plot.

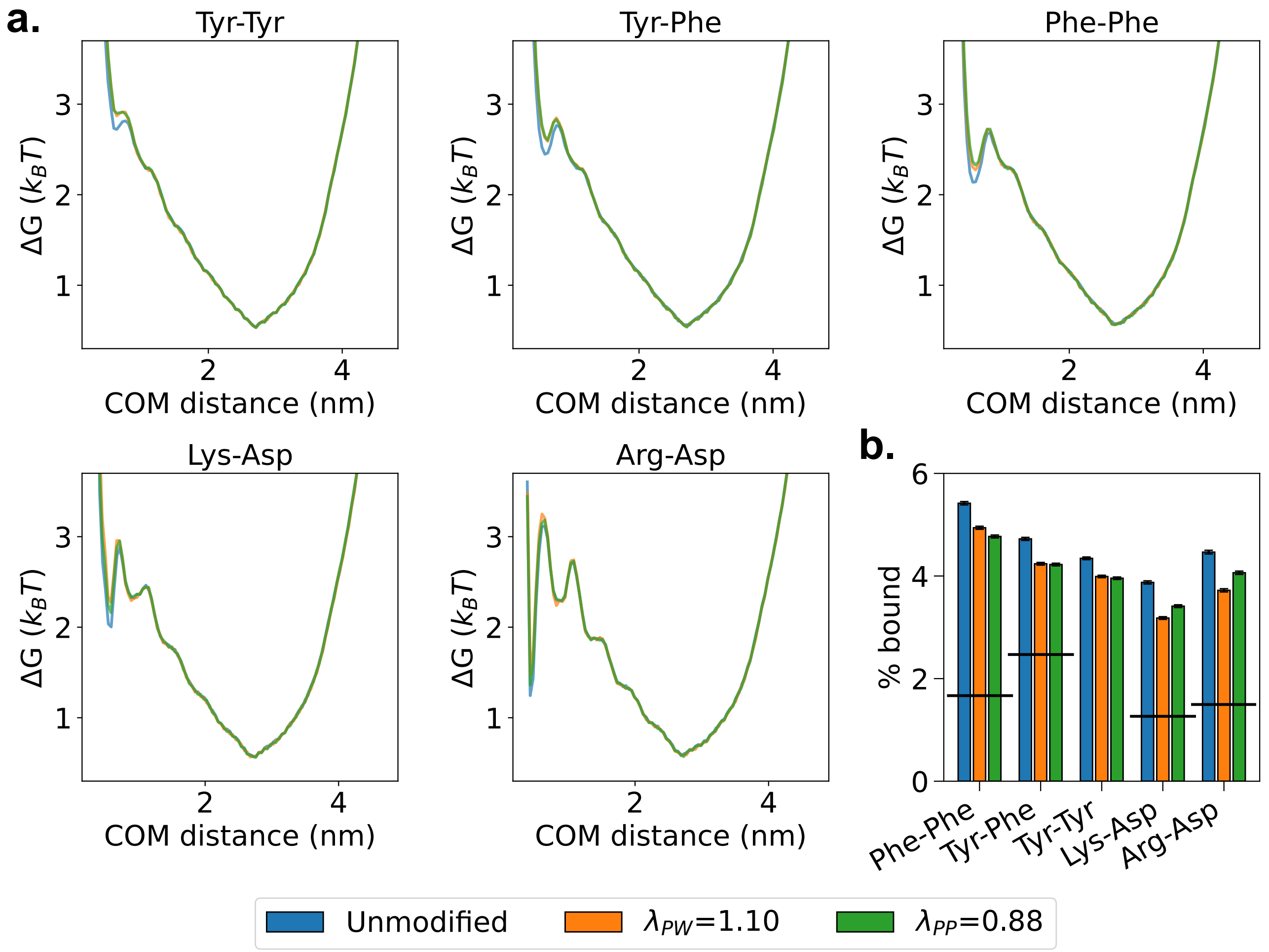


**Figure S9. Self-association of amino acid side chain analogs.** Simulations of the self-association of amino acid side chain analogs with unmodified Martini 3 (blue), protein-water interactions in Martini 3 rescaled by $\lambda_{PW}$=1.10 (orange), and protein-protein interactions in Martini 3 rescaled by $\lambda_{PP}$=0.88 (green). **a.** Free energy profile along the center-of-mass (COM) distance between the two side chain analogs calculated from simulations with the three different force fields. **b.** Fraction bound calculated from simulations with the three different force fields. The black lines indicate the expected fraction bound given by experimental $K_a$-values of 0.4 $M^{-1}$ for Phe-Phe (benzene-benzene), 0.6 $M^{-1}$ for Phe-Tyr (benzene-phenol) (***Christian and Tucker, 1982***), 0.31 $M^{-1}$ for Lys-Asp (butylammonium-acetate), and 0.37 $M^{-1}$ for Arg-Asp (guanidine-acetate) (***Springs and Haake, 1977***).

| Protein | $N_R$ | Domain 1 | Domain 2 | Domain 3 | Domain 4 |
|---|---|---|---|---|---|
| MyBP-$C_{MTHB\text{-}C2}$ | 137 | 6-42 | 50-137 | - | - |
| $Ubq_2$ | 162 | 11-82 | 87-158 | - | - |
| $Ubq_3$ | 228 | 1-72 | 77-148 | 153-224 | - |
| Gal-3 | 250 | 117-250 | - | - | - |
| TIA1 | 275 | 6-82 | 95-172 | 190-275 | - |
| $Ubq_4$ | 304 | 1-72 | 77-148 | 153-224 | 229-300 |
| hnRNPA1 | 314 | 11-89 | 105-179 | - | - |
| MyBP-$C_{C5\text{-}C6\text{-}C7}$ | 328 | 8-133 (8-51) (89-133) | 142-215 | 236-328 | - |
| PTEN | 351 | 1-184 | 190-351 | - | - |
| Talin FERM | 368 | 1-83 | 85-195 | 208-304 | 311-368 |
| hisSUMO-hnRNPA1 | 433 | 44-114 | 132-209 | 224-298 | - |
| mTurq-mNeon | 470 | 1-226 | 256-470 | - | - |
| mTurq-$GS_8$-mNeon | 486 | 1-226 | 272-486 | - | - |
| mTurq-$GS_{16}$-mNeon | 502 | 1-226 | 288-502 | - | - |
| mTurq-$GS_{24}$-mNeon | 518 | 1-226 | 304-518 | - | - |
| mTurq-$GS_{32}$-mNeon | 534 | 1-226 | 320-534 | - | - |
| mTurq-$GS_{48}$-mNeon | 566 | 1-226 | 352-566 | - | - |

**Table S1. Domain boundaries for assignment of the elastic network model.** Shown are the total number of amino acid residues ($N_R$) and the residues within the specified domain boundaries (domain 1, 2, 3, and 4). For MyBP-$C_{C5\text{-}C6\text{-}C7}$, elastic network restraints were removed from the loop region (res. 52–88) of the C5 domain (domain 1).

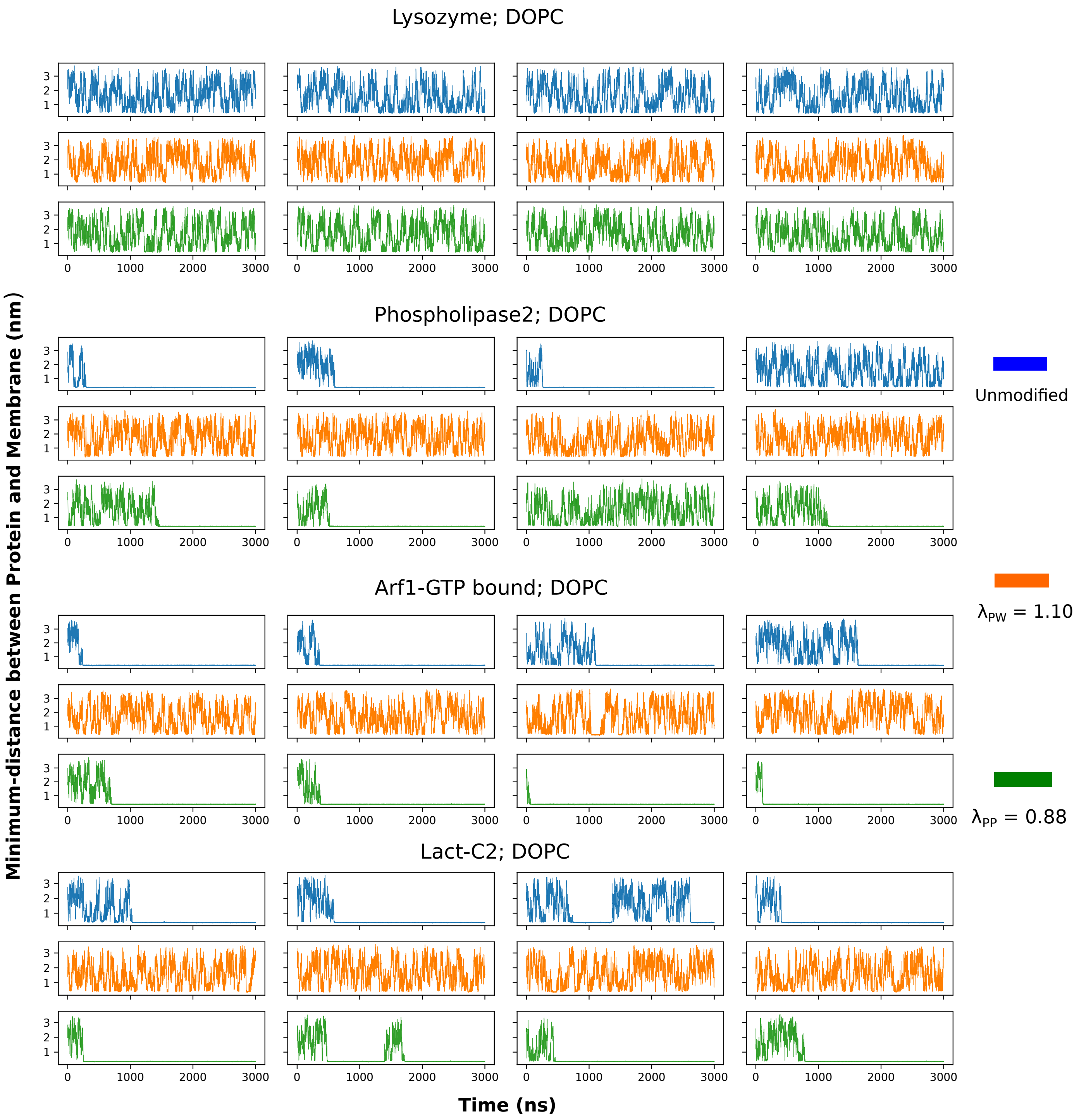


**Figure S10.** Time evolution of the minimum distance between protein and membrane for three peripheral membrane proteins and one negative control (Lysozyme) from simulations with unmodified Martini 3 (blue), protein-water interactions in Martini 3 rescaled by $\lambda_{PW}$=1.10 (orange), and protein-protein interactions in Martini 3 rescaled by $\lambda_{PP}$=0.88 (green). Four replicas of each system and parameter combination are shown. We note that the slow timescale of membrane dissociation means that it is in some cases difficult to obtain a converged value for the fraction of time bound to the membrane.

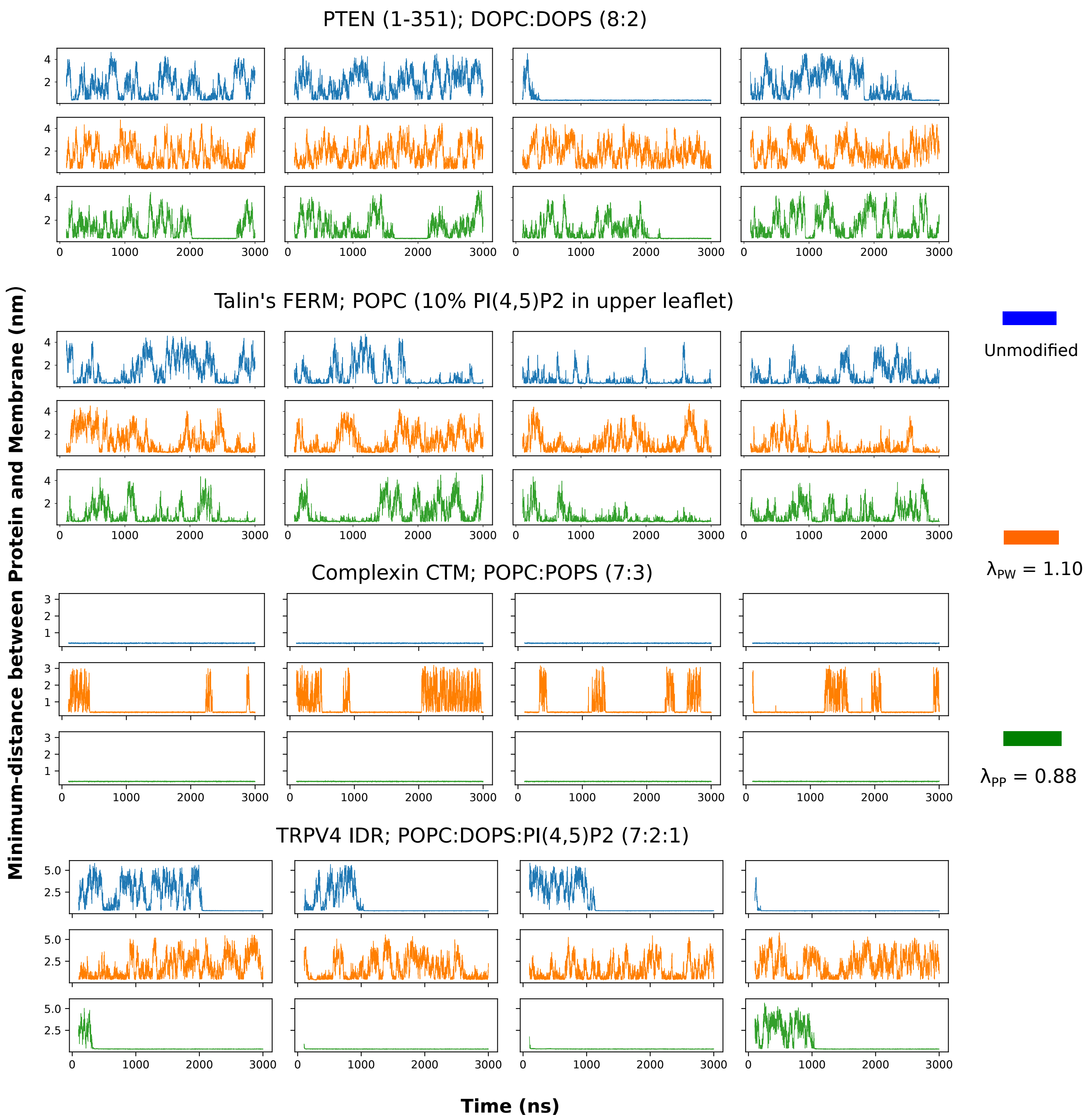


**Figure S11.** Time evolution of the minimum distance between protein and membrane for two multidomain proteins and two IDPs from simulations with unmodified Martini 3 (blue), protein-water interactions in Martini 3 rescaled by $\lambda_{PW}$=1.10 (orange), and protein-protein interactions in Martini 3 rescaled by $\lambda_{PP}$=0.88 (green). Four replicas of each system and parameter combination are represented. We note that the slow timescale of membrane dissociation means that it is in some cases difficult to obtain a converged value for the fraction of time bound to the membrane.

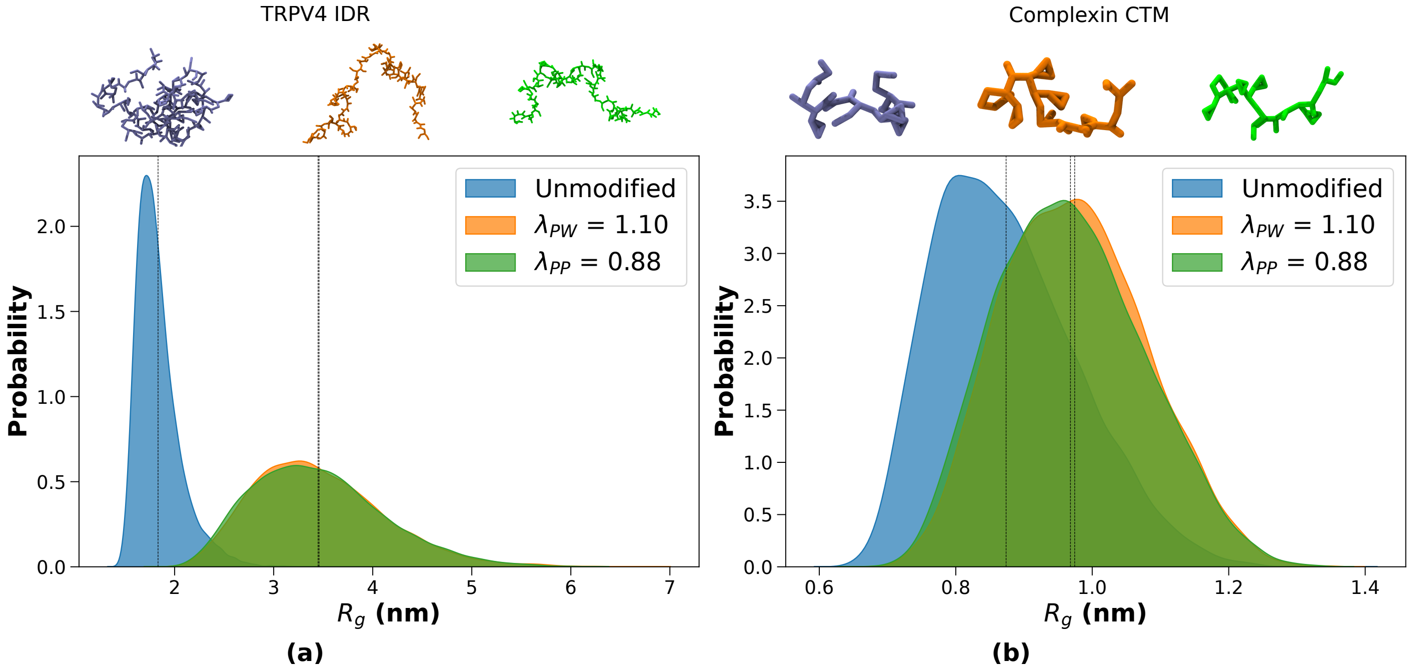


**Figure S12.** Probability distributions of the radius of gyration ($R_g$) for (a) TRPV4 IDR, and (b) Complexin CTM from simulations with unmodified Martini 3 (blue), protein-water interactions in Martini 3 rescaled by $\lambda_{\rm PW}$=1.10 (orange), and protein-protein interactions in Martini 3 rescaled by $\lambda_{\rm PP}$=0.88 (green). Black dashed lines represent the average values of $R_g$.
